# Comprehensive analysis of pyroptosis-associated in molecular classification, immunity and prognostic of glioma

**DOI:** 10.1101/2021.08.03.454997

**Authors:** Peng Chen, Yanyan Li, Na Li, Liangfang Shen, Zhanzhan Li

## Abstract

Integrative analysis was performed in the Chinese Glioma Genome Atlas and The Cancer Genome Atlas to describe the pyroptosis-associated molecular classification and prognostic signature in glioma. Pyroptosis-related genes were used for consensus clustering and to develop a prognostic signature. The immune statuses, molecular alterations and clinical features of differentially expressed genes were analyzed among different subclasses and risk groups. A lncRNA-miRNA-mRNA network was built, and drug sensitivity analysis was used to identify small molecular drugs for the identified genes. Glioma can be divided into two subclasses using 30 pyroptosis-related genes. Cluster 1 displayed high immune signatures and poor prognosis as well as high immune-related function scores. A prognostic signature based on 15 pyroptosis-related genes of the CGGA cohort can predict the overall survival of glioma and was well validated in the TCGA cohort. Cluster 1 had higher risk scores. The high-risk group had high immune cell and function scores and low DNA methylation of pyroptosis-related genes. The differences in pyroptosis-related gene mutations and somatic copy numbers were significant between the high-risk and low-risk groups. The ceRNA regulatory network uncovered the regulatory patterns of different risk groups in glioma. Nine pairs of target genes and drugs were identified. In vitro, CASP8 promotes the progression of glioma cells. Pyroptosis-related genes can reflect the molecular biological and clinical features of glioma subclasses. The established prognostic signature can predict prognosis and distinguish molecular alterations in glioma patients. Our comprehensive analyses provide valuable guidelines for improving glioma patient management and individualized therapy.

## INTRODUCTION

Gliomas are the most common types of primary tumors in the central nervous system and one of the most devastating tumors (1). At present, the main treatment methods of glioma are surgical resection, radiotherapy, chemotherapy or chemoradiotherapy(2). Although great efforts have been made to improve glioma treatment, the prognosis of glioma patients remains poor(3). One of the main reasons is that the molecular mechanism is still not fully understood. Therefore, the exploration and research of the underlying mechanism of gliomas and identification of potential treatment targets followed by application in clinical practice have important theoretical and practical significance.

Pyroptosis is one of the pathways involved in programmed cell death, such as apoptosis, ferroptosis, necroptosis, and autophagy.(4) Cookson et al. first used pyroptosis to describe the caspase-1-dependent pattern of cell death found in macrophages(5). Pyroptosis, distinct from apoptosis and necrosis, contributes to a range of human diseases as a new mechanism of cell death. Pyroptosis is a proinflammatory form of programmed cell death that is dependent on the activity of caspase acid-specific proteases(6). In the coupling of the amino-terminal and carboxy-terminal linkers of gasdermin D (GSDMD) by caspases, the latter is displaced onto the membrane and perforated, inducing moisture penetration, cell swelling and the release of inflammatory factors, which is followed by pyroptosis(7). A previous study reported that pyroptosis plays an important role in immunity and diseases. Pyroptosis can promote the death of damaged cells during infection and acts as an alarm signal for the recruitment of immune cells to the site of infection to promote the removal of pathogens, thus effectively protecting the body(8). In recent years, its role in tumorigenesis and cancer development has been studied comprehensively. Various regulators have been reported to be involved in the process of pyroptosis and play pivotal roles in the progression of tumors, such as hepatocellular carcinoma, lung cancer, and breast cancer(9-11). However, few studies have investigated the role of pyroptosis in glioma, and comprehensive analyses of pyroptosis regulators in glioma, their correlation with clinical characteristics and their prognostic value have not been reported.

In the present study, we first outlined the molecular subtypes of gliomas based on pyroptosis-related genes in the CGGA dataset and described the clinical and molecular characteristics and immune status of each subclass. Then, we developed a prognostic signature of pyroptosis-related genes based on the CGGA cohort, validated this prognostic signature in the TCGA cohort. Furthermore, we explored the clinical and molecular patterns, including immune infiltration, somatic copy number alterations, mutations, and DNA methylation, and established a lncRNA-miRNA-mRNA regulatory network. Finally, we explored the correlation between small molecular drugs and the identified prognostic signature genes. Our comprehensive analyses provide new insight into the functions of pyroptosis in the initiation, development, and progression of glioma.

## MATERIALS AND METHODS

### Data source

We downloaded the genomic data, copy number alteration, methylation and clinical data of glioma patients from the CGGA (http://www.cgga.org.cn/) and TCGA databases (https://portal.gdc.cancer.gov/). Additional gene-centric RMA-normalized gene expression profiles and drug response data of over 1000 cancer cell lines were accessed from the Genomics of Drug Sensitivity in Cancer (GDSC) database (https://www.cancerrxgene.org/downloads). Immune-associated data, including immune cells and immunophenoscores, were downloaded from TCIA (https://tcia.at/home). Thirty-three pyroptosis-related genes were defined from a previous publication and are provided in Table S1(12-15).

### Identification of glioma subclasses and Gene set variation analysis

We identified the optimal clustering number visualizing consensus matrix, tracking plot, and cumulative distribution function plot. In addition, a T-distributed stochastic neighbor embedding-based approach was used to validate the clustering in glioma patients. We calculated the enrichment scores for every sample using the GSVA R package.

### Development and validation of a prognostic signature

We developed a pyroptosis-related prognostic signature based on the CGGA training cohort. Twenty differentially expressed genes with P<0.05 were entered into LASSO Cox regression, which identified potential genes for the prognostic signature in the CGGA training cohort. Then, we calculated the risk score for each sample of the CGGA and TCGA validation cohorts using the obtained regression coefficient in the CGGA training cohort: risk score =coef_1_*gene_1_ expression+coef_2_* gene_2_ expression+…coefn*genen expression. The CGGA and TCGA samples were divided into a high-risk group and a low-risk group based on the median risk score. Receiver operating characteristic curves were plotted to evaluate the 1-year, 2-year, and 3-year sensitivity and specificity of the prognostic signature. We also established a prognostic nomogram to evaluate the clinical value of the prognostic signature. Calibration analysis of the prognostic predictive value of the nomogram was carried out.

### Functional enrichment analysis, estimation of tumor stem cell-like properties and immune infiltration

Gene Ontology and KEGG pathway analyses were performed using the “clusterProfiler” package. We used single-sample gene set enrichment analysis (ssGSEA) to estimate the enrichment score of stem cell-like properties (RNAss, DNAss) and the TME (stromal score, immune score, and ESTIMATE score) in the TCGA cohort because the CGGA dataset did not provide such data. The immune-related cell and function scores were also calculated for each sample (downloaded from https://www.gsea-msigdb.org/).

### Somatic copy number alteration, mutation, and DNA methylation analysis

Based on the risk groups in the TCGA cohort, we compared the somatic copy number alteration, mutation, and DNA methylation levels between the high-risk and low-risk groups using the “limma” R package.

### Construction of a ceRNA network and drug sensitivity

To further explore the transcriptome regulation network of different risk groups, we used Cytoscape version 3.8.2 to establish a lncRNA-miRNA-mRNA regulatory network. We explore the correlation between small molecular drugs and the identified prognostic signature genes using Pearson correlation analysis |R|>0.25 and P>0.05 were considered significant.

### Verification of experiments in vitro

We further performed the Western blot, cell migration assays, cell scratchy assays, and clonogenic assays to verify the present finding. We selected the CASP8 to validate the molecular function because CASP8 showed significant differences between normal tissue and GBM or LGG, and the elevated expression is associated with poor prognosis. The details of experiments process in vitro were supplied in Additional file 1.docx.

### Statistical analysis

The log-rank test was used to compare the survival curves of Kaplan-Meier analysis. The hazard ratio (HR) and 95% confidence interval (CI) of each gene and clinical parameters were calculated when univariate and multivariate Cox regression were applied. All analyses were achieved using R software version 4.0. A two-sided P value <0.05 was considered significant unless otherwise specified.

## RESULTS

### Identification of glioma subclasses

The flow chart of the data analysis is presented in Figure 1A. From two CGGA RNA-seq datasets, we obtained 1018 samples of gene expression data and further identified 30 pyroptosis-related genes based on MAD>0.5. The gene symbols and descriptions of the 30 pyroptosis-associated genes used for classification are listed in the Additional file 2: Table S1. We first explored the interactions among these genes using PPIs (Figure 1B), and the PPI network indicated that CASP8, CASP4, CASP1, NLRP3, NLRP1 and NLRC4 are hub genes. The correlation circle plot of the 30 genes is presented in Figure 1C (red: positive correlation; green: negative correlation). We identified the optimal k value as 2 by estimating the comprehensive correlation coefficient. Therefore, we divided the glioma samples into two different subclasses: cluster 1 and cluster 2. For the optimal k value (k=2), the consensus matrix showed a relatively sharp and clear boundary, indicating stable and robust clustering (Figure 1D). To verify the subclass stability, we further performed t-sensitivity PCA and found that a two-dimensional t-sensitivity distribution supported subtype clustering (Figure 1E). The consensus clustering for each sample is listed in Additional file 2: Table S2. The Kaplan-Meier analysis indicates that the median survival time was significantly shorter in cluster 2 than in cluster 1 (MST: 1.87 vs. 6.92 years, P<0.001, Figure 1F). This result indicated that the two subclasses had distinct prognostic patterns.

**Figure 1.**
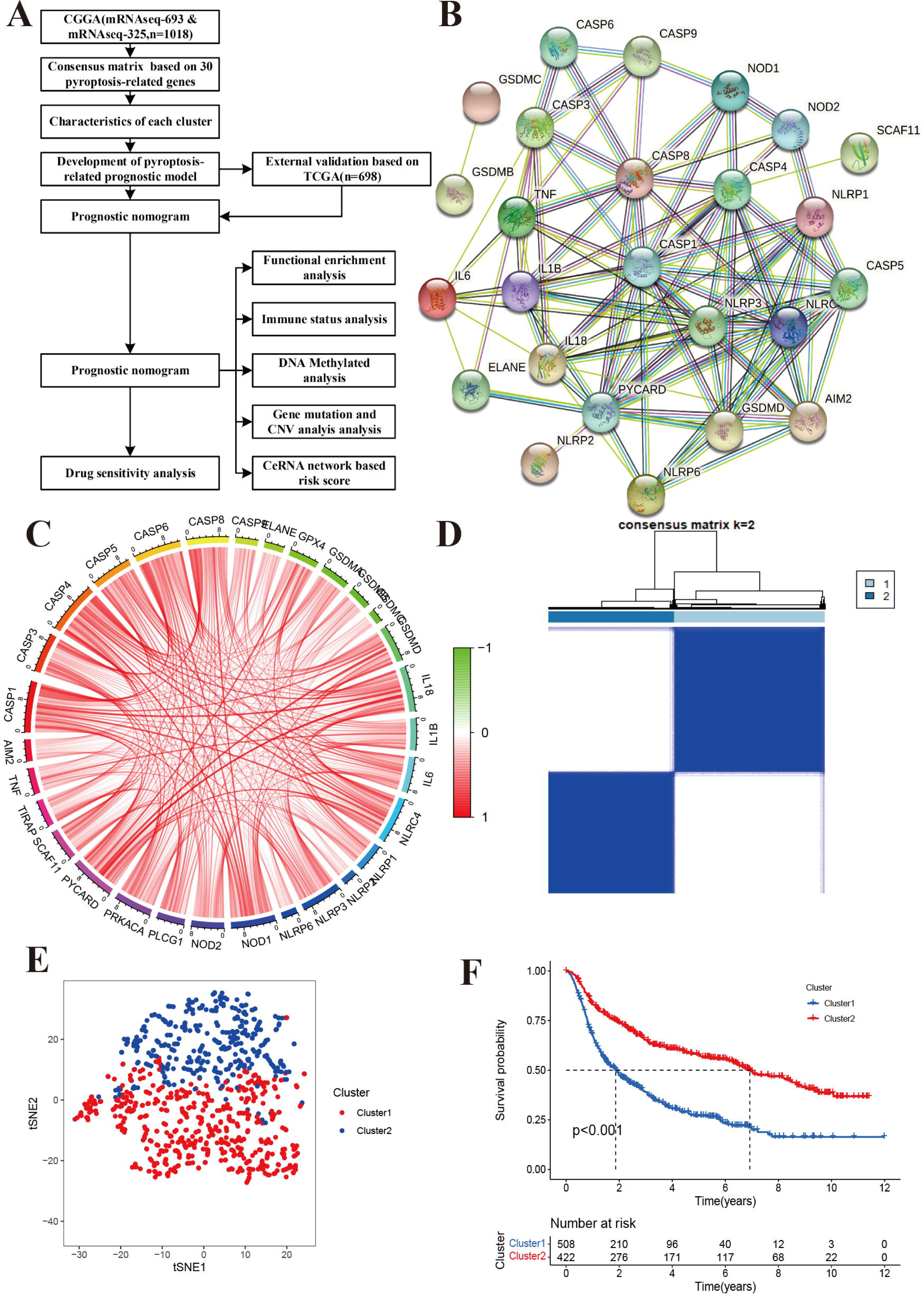
Identification of glioma subclasses using consensus clustering method in the CGGA dataset. **(A)** Flow chart of the study. **(B)** PPI network indicating the interactions among pyroptosis-related genes (interaction score=0.7). **(C)** The circle plot of correlation among pyroptosis-related genes (green line: negative correlation, red line: positive correlation). **(D)** Consensus matrix method clustering using 30 pyroptosis-related genes. **(E)** PCA analysis showed the distribution of two glioma subclasses in the CGGA dataset. **(F)** Overall survival curve of two clusters in the cohort.

### Correlation of glioma subclasses with pyroptosis-related genes

Two subclasses were obtained based on pyroptosis-related genes. To explore the pathway enrichment for the two subclasses, we performed GSVA by transforming the expression data from a gene-by-sample matrix to a gene set by two subclasses. Then, differential pathways were enriched in the two subclasses. Compared with cluster 1, the GSVA results indicated that cluster 2 had 182 kinds of significantly differential signaling pathways (Additional file 2: Table S3). The upregulated pathways were associated with immune-related pathways, such as autoimmune, allograft rejection, graft versus host disease, primary immunodeficiency, antigen processing and presentation. Some signaling pathways, such as the cytosolic DNA sensing pathway, NOD-like receptor signaling pathway, Toll-like receptor signaling pathway, and metabolism-related pathways, were also significantly enriched. The significantly downregulated pathway was long-term potentiation (Figure 2A).

**Figure 2.**
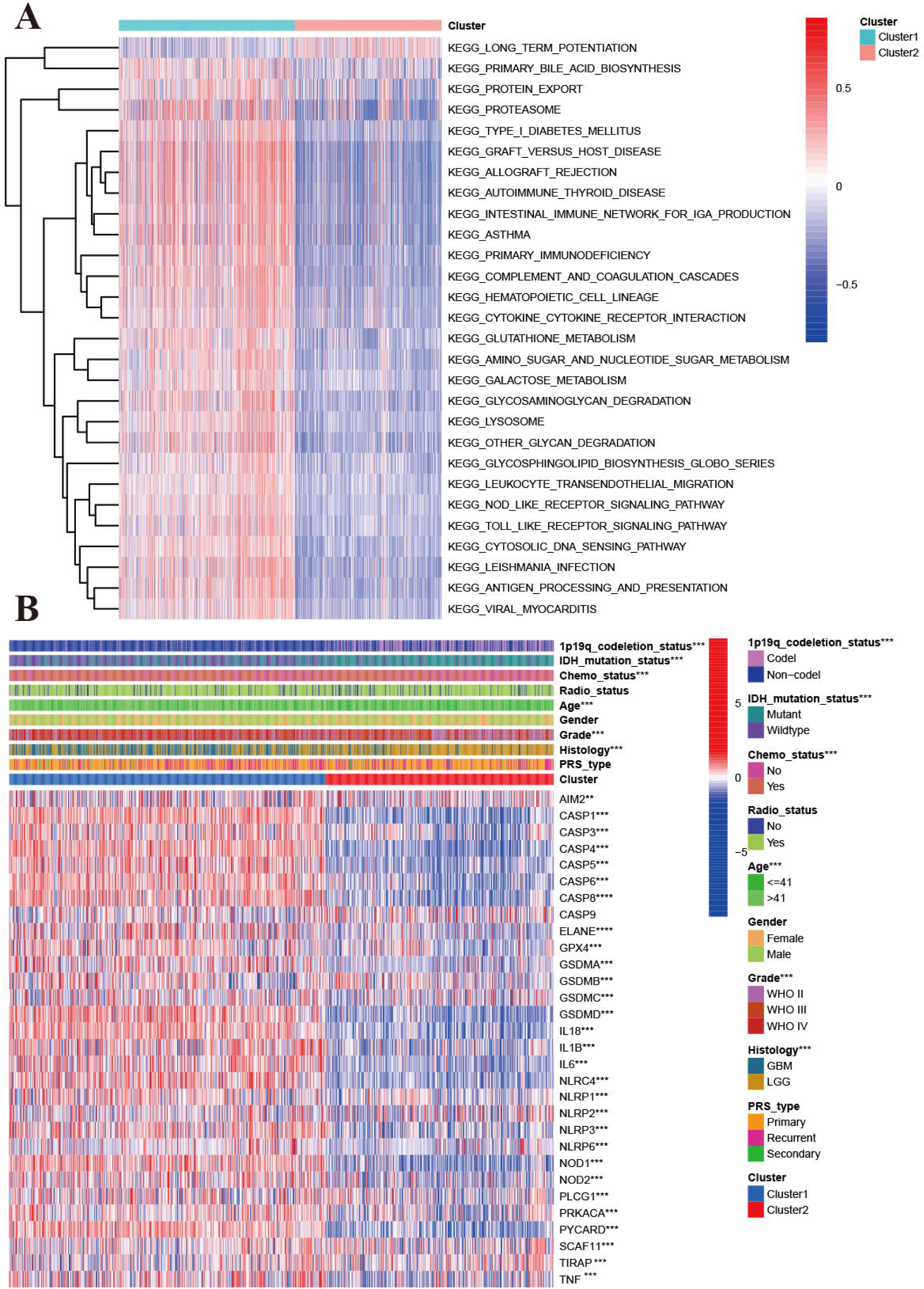
Characteristics of patients in cluster 1 and cluster 2 in CGGA cohort. **(A)** Heatmap of gene set variation analysis of the pyroptosis-related genes from cluster 1 and cluster 2. **(B)** Heatmap showed the correlations between two subclasses and clinical characteristics and differentially expressed pyroptosis-related genes in the CGGA cohort.

### Clinical characteristics and transcriptomes of glioma subclasses

We explored the correlation of subclasses with clinical characteristics (Figure 2B). Compared with patients in cluster 2 with a favorable prognosis, patients in cluster 1 tended to have GBM (P<0.001), WHO grade IV (P<0.001), a higher proportion of age >41 years, 1p19q non-codeletion status (P<0.001), and IDH wildtype status (P<0.001). Sex, PRS type and radiotherapy status were not associated with the molecular subclasses (P>0.05). For the pyroptosis-related genes except CASP9, significant differential expression was observed in the two clusters. Among these differentially expressed genes, all genes were upregulated in cluster 1 and downregulated in cluster 2 (Figure 2B). We also compared the differences in pyroptosis-related genes in patients with different histologies, grades, IDH mutation statuses, and 1p19q statuses. Compared with the LGG group, the GBM group had one upregulated gene (AIM2) and 21 downregulated genes (Additional file 3: Figure S1A). Twenty-one DEGs were found for grade, and their expression increased with increasing WHO grade (P<0.005, Additional file 3: Figure S1B). For IDH status, 25 DEGs were found (Additional file 3: Figure S1C). Thirty pyroptosis-related DEGs were found for 1p191 status (Additional file 3: Figure S1D).

We further performed differential expression analysis between cluster 1 and cluster 2. A total of 392 DEGs were found, 18 genes were upregulated, and 372 genes were downregulated in cluster 2 (Additional file 2: Table S4). GO and KEGG enrichment analyses were performed for all DEGs (Additional file 2: Table S5 and Table S6). A total of 874 differentially expressed functions were enriched, including 709 biological processes, 95 cellular components and 70 molecular functions. The top 30 enrichment results are presented in Additional file 3: Figure S2. Most of these functions were associated with immunity. In addition, 56 pathways were also identified in the KEGG analysis (Additional file 3: Figure S3), and the top five pathways were phagosome, Staphylococcus aureus infection, tuberculosis, complement and coagulation cascades, and human T-cell leukemia virus 1 infection.

### Correlation of glioma subclasses with immune status

To explore the tumor heterogeneity between the two subclasses, we investigated the immune cell and immune function differences. Compared with cluster 2, cluster 1 had higher aDC, CD8+ T cell, DC, iDC, macrophage, mast cell, neutrophil, NK cell, pDC, T helper cell, Tfh cell, Th2 cell, TIL, and Treg levels (all P<0.001, Additional file 3: Figure S4A). Similarly, cluster 1 had higher immune function scores than cluster 2, including APC coinhibition, APC costimulation, CCR, checkpoint, cytolytic activity, HLA, inflammation promotion, MHC class I, parainflammation, T cell coinhibition, type I IFN response and type II IFN response (all P<0.001, Additional file 3: Figure S4B).

### Development of a pyroptosis-related prognostic signature in glioma

Initially, we performed univariate Cox regression to identify the correlations of the 30 pyroptosis-related genes with OS (Additional file 3: Figure S5A) in the CGGA cohort. In total, 20 pyroptosis-related genes were identified as associated with the overall survival of glioma patients. The Kaplan-Meier plot indicated that high expression of CASP3, CASP4, CASP5, CASP6, CASP8, ELANE, GSMAD, IL6, NLRP3, NOD1, NOD2, PLCG1, PRKACA, PYCARD, and SCAF11 was associated with poorer OS in glioma. Using 20 prognostic pyroptosis-related genes, we developed a prognostic signature by performing LASSO regression in the CGGA training cohort (Additional file 3: Figure S5B and S5C). Fifteen of the 20 prognostic genes were used to develop the risk signature. We calculated the risk score for each sample using the regression coefficients of the 15 genes (Additional file 2: Table S7). Glioma patients with risk scores greater than the median value were divided into a high-risk group, and the others were divided into a low-risk group. Compared with the low-risk group, the high-risk group was more likely to have GBM (P<0.001), a higher WHO grade (P<0.001), recurrence (P<0.001), older age (P<0.001), IDH wildtype status (P<0.001), 1p19q non-codeletion status (P<0.001), and a history of chemotherapy (P<0.001). The heatmap showed the association between the risk group and clinical parameters and differentially expressed genes of the high- and low-risk groups (Additional file 3: Figure S5D). Furthermore, we found that glioma patients belonging to cluster 1, patients with a poor prognosis, patients with GBM, patients with WHO grade IV patients with 1p19q non-codeletion status and patients with IDH wildtype status had higher risk scores (all P<0.001, Additional file 3: Figure S6).

The Kaplan-Meier analysis showed that the high-risk group had a significantly poorer OS than the low-risk group (Figure 3A-3 B). Univariate Cox regression indicated that the risk score was positively associated with OS in glioma (HR=3. 105, 95% CI: 2.681– 3.596, P<0.001, Figure 3C). Multivariate Cox regression suggested that the risk score was an independent unfavorable prognostic predictor in glioma (HR=1.685, 95% CI: 1.392–2.039, P<0.001, Figure 3D). In addition, PRS type, tumor grade, and age were positively associated with OS. However, chemotherapy, wildtype IDH status, and 1p19q status were negatively associated with OS in the CGGA training cohort. The PCA plot indicated that patients in different risk groups were separated into obviously different clusters (Figure 3E). Time-dependent receiver operating characteristic analysis was performed to evaluate the predictability of the prognostic model. Our results showed that the AUCs at 1 year, 2 years, and 3 years were 0.717, 0.784 and 0.773 (Figure 3F), respectively. We further compared the OS status among different histology, IDH status, 1p19q codeletion status, and grade subgroups. The results showed that the OS of the high-risk group was still poorer than that of the low-risk group (Additional file 3: Figure S7, all P<0.001).

**Figure 3.**
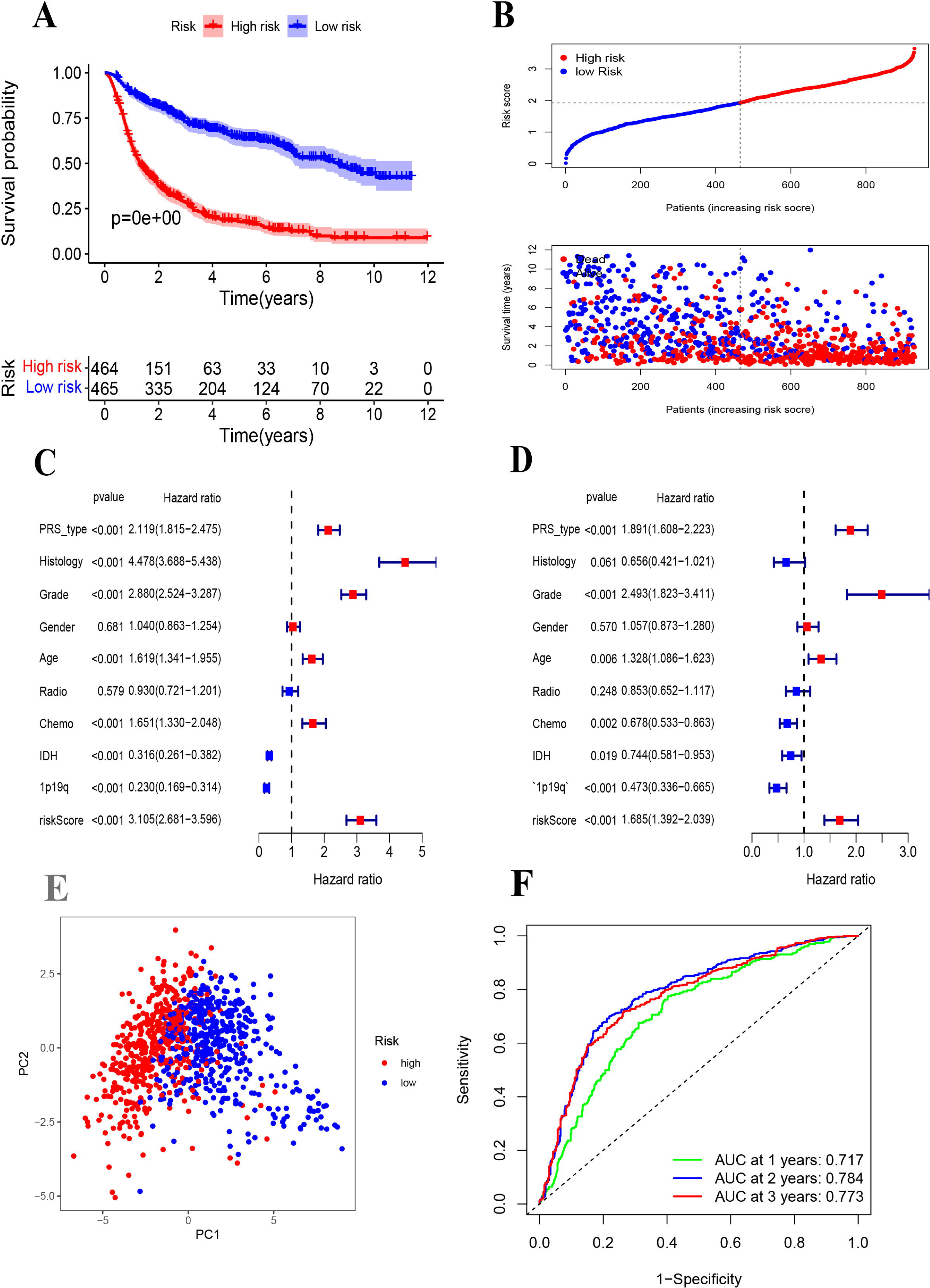
Establishment of a pyroptosis-related gene prognostic signature in the CGGA cohort. **(A)** Kaplan-Meier curves for OS of patients in high- and low-risk group in CGGA Cohort. **(B)** Distribution of risk score of all patients of CGGA cohort, and Patients’ survival time distribution. **(C)** Forest plot of univariate cox regression between risk score and prognosis of glioma. **(D)** Forest plot of multivariate cox regression of between risk score and prognosis of glioma. **(E)** PCA plot for signature genes based on risk score group. **(F)** ROC curves showed the predictive efficiency of risk score at 1-year, 2-year, 3-year point.

### External validation of the pyroptosis-related prognostic signature in glioma

To further validate the prognostic value of the pyroptosis-related gene model, we also calculated the risk score of glioma patients in the TCGA cohort using the regression coefficients of the CGGA cohort. The Kaplan-Meier analysis indicated a significant correlation of the high-risk group with worse OS than the low-risk group (Figure 4A-4C). Univariate Cox regression showed that the risk score was significantly associated with OS in the TCGA cohort (HR=2.084, 95% CI: 1.890-2.297, P<0.001, Figure 4D).

**Figure 4.**
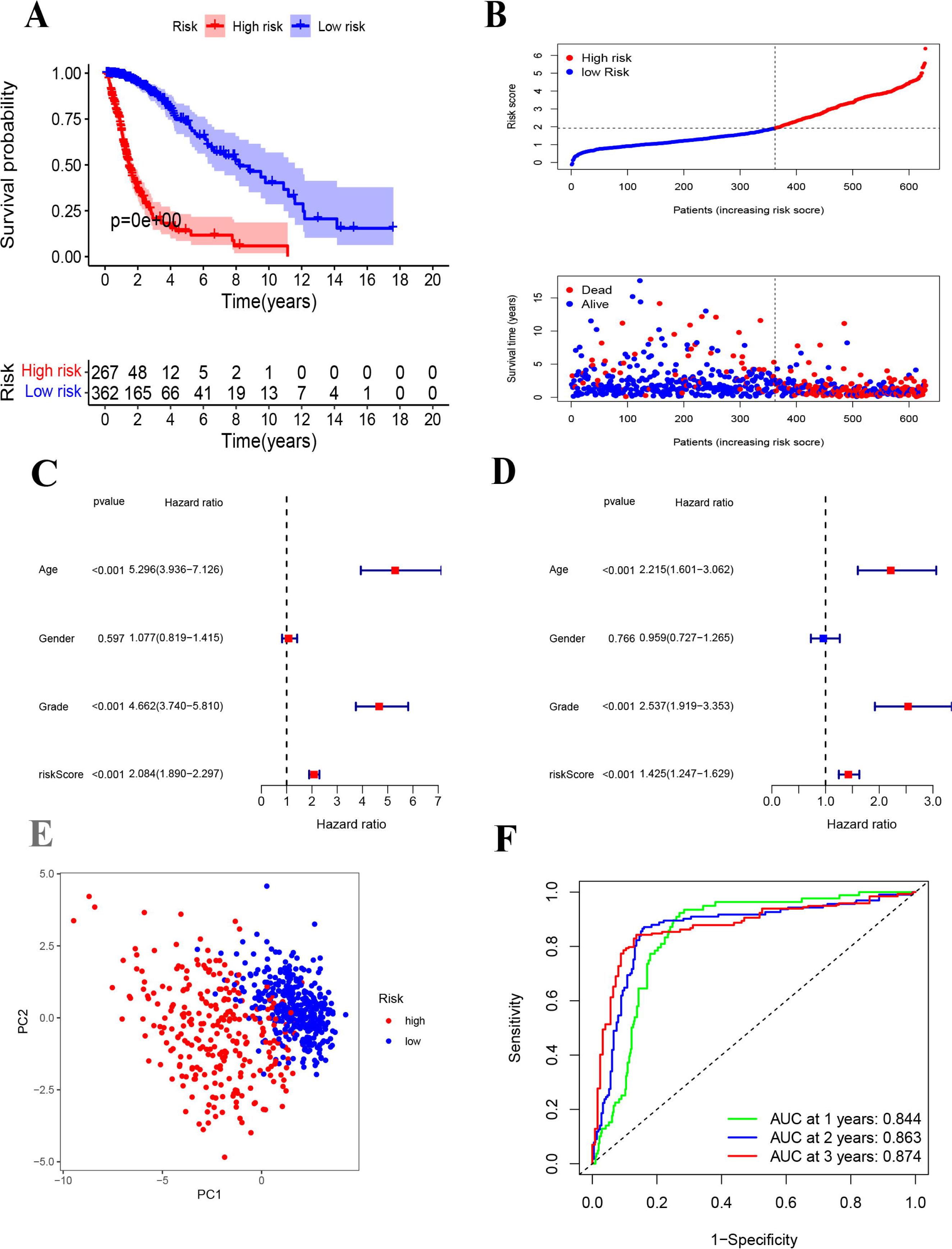
External validation of a pyroptosis-related gene prognostic signature in the TCGA cohort. **(A)** Kaplan-Meier curves for OS of patients in high- and low-risk group in TCGA Cohort. **(B)** Distribution of risk score of all patients of TCGA cohort and Patients’ survival time distribution of TCGA cohort. **(C)** Forest plot of univariate cox regression between risk score and prognosis of glioma in TCGA cohort. **(D)** Forest plot of multivariate cox regression of between risk score and prognosis of glioma in TCGA cohort. **(E)** PCA plot for signature genes based on risk score group in TCGA cohort. **(F)** ROC curves showed the predictive efficiency of risk score at 1-year, 2-year, 3-year point in TCGA cohort.

In multivariate Cox regression, the risk score was also an independent prognostic indicator (HR=1.425, 95% CI: 1.247–1.629 P<0.001, Figure 4E). The PCA plot validated the high- and low-risk distribution of all glioma patients based on the TCGA cohort. Furthermore, the AUCs of the risk score were 0.844 at 1 year, 0.863 at 2 years, and 0.874 at 3 years (Figure 4F).

### Prognostic prediction models

To further evaluate the clinical prediction value of the prognostic signature, we constructed a prognostic nomogram model based on multivariate Cox regression analysis that included all clinical parameters in the CGGA cohort The calibration curves indicated that the clinical nomogram model could precisely predict the 1-year, 3-year and 5-year OS of glioma patients (C-index=0.799). The predictive accuracy of this nomogram was well validated in the TCGA cohort (C-index=0.841, Figure 5).

**Figure 5.**
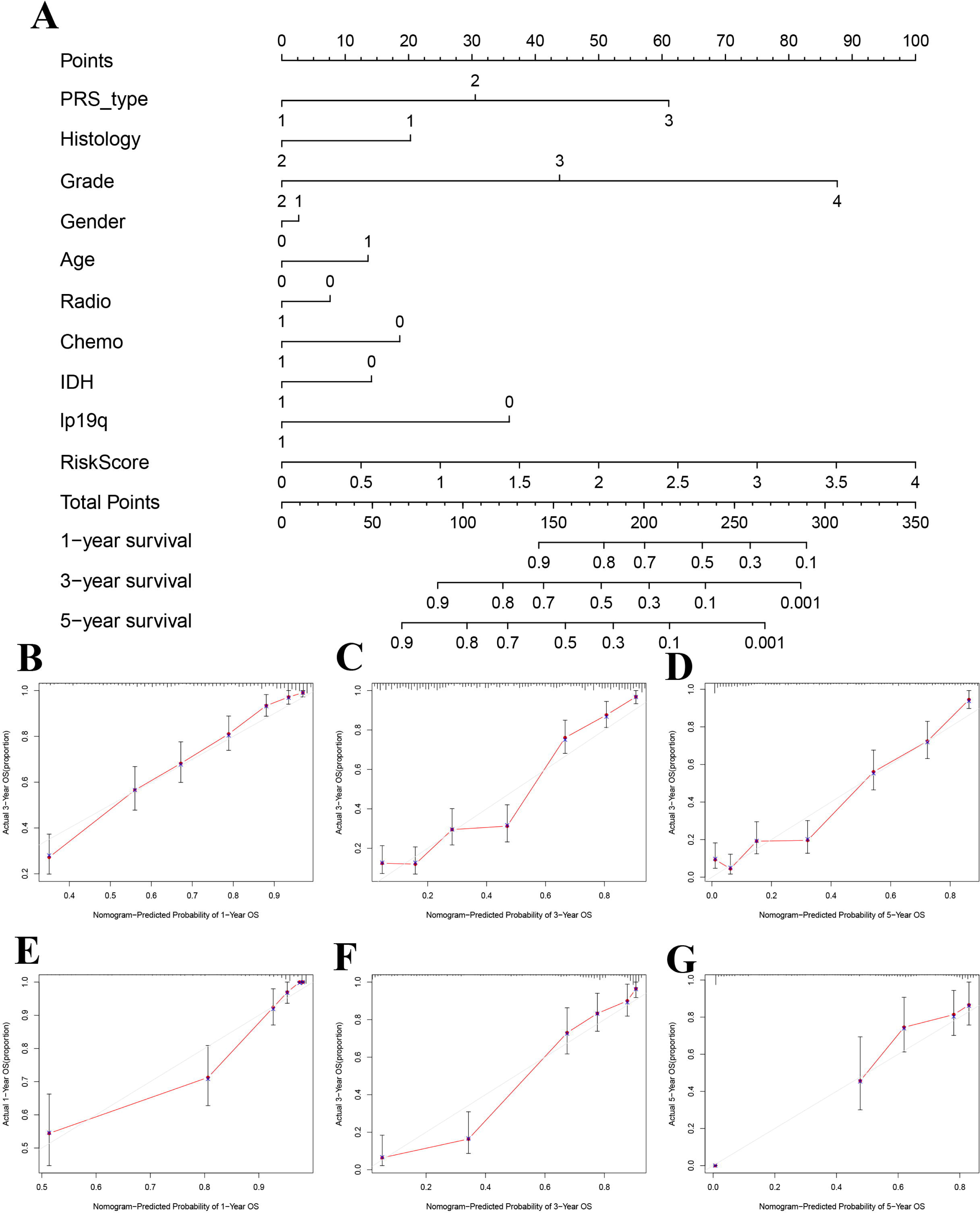
Establishment and validation of nomogram model based on prognostic signature genes. (A) Nomogram model established in the CGGA cohort. (B) The 1-year calibration curves in the CGGA cohort. (C) The 3-year calibration curves in the CGGA cohort. (D) The 5-year calibration curves in the CGGA cohort. (E) The 1-year calibration curves in the TCGA cohort. (F) The 3-year calibration curves in the TCGA cohort. (G) The 5-year calibration curves in the TCGA cohort.

### Functional enrichment and immune infiltration analyses based on the prognostic signature

We further explored the underlying biological functions that define the survival of glioma patients. We first performed DEG analysis between the high-risk and low-risk groups and then annotated the functions of the DEGs in terms of biological processes, cellular components, and molecular functions using GO enrichment and KEGG pathways. We identified 338 DEGs in the CGGA cohort (Additional file 2: Table S8) and 2600 DEGs in the TCGA cohort (Additional file 2: Table S9). The GO enrichment and KEGG pathway analyses indicated that the CGGA and TCGA cohorts shared some enrichment results, such as extracellular matrix organization, extracellular structure organization, immune response, ECM-receptor interaction, and cell adhesion molecules (Figure 6A-6D).

**Figure 6.**
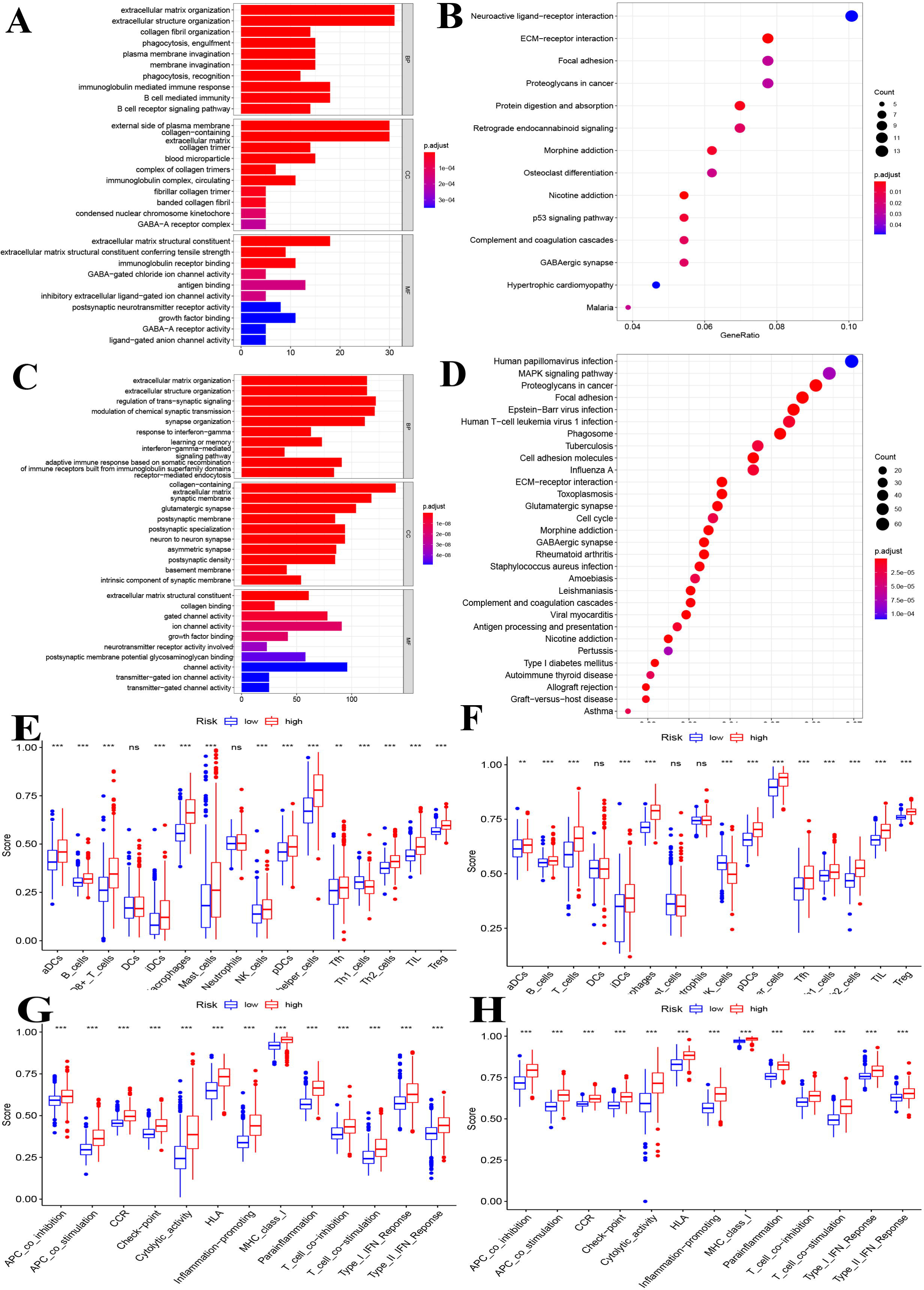
Functional enrichment and immune status analysis. **(A)** Barplot of enrichment analysis based on prognostic-related signature genes in CGGA cohort. **(B)** Bubble plot of enrichment analysis based on prognostic-related signature genes in CGGA cohort. **(C)** Barplot of enrichment analysis based on prognostic-related signature genes in TCGA cohort. **(D)** Bubble plot of enrichment analysis based on prognostic-related signature genes in TCGA cohort. **(E)** Boxplot showed the ssGSEA scores for immune cells based on risk group in CGGA cohort. **(F)** Boxplot showed the ssGSEA scores for immune cells based on risk group in TCGA cohort. **(G)** Boxplot showed the ssGSEA scores for immune pathways based on risk group in CGGA cohort. **(H)** Boxplot showed the ssGSEA scores for immune pathways based on risk group in TCGA cohort

We also explored the differences in immune cells and immune functions based on the risk score in the CGGA (Figure 6E and Figure 6G) and TCGA datasets (Figure 6F and Figure 6H). As shown in the box plots, the immune cell score showed a similar trend in the CGGA and TCGA datasets. All immune cell scores were significantly upregulated in the high-risk group. The immune function differences of the different risk groups were the same in the CGGA and TCGA datasets (all P<0.001). All immune function scores were significantly upregulated in the high-risk group. Significant expression levels were also observed among different immune subtypes, which indicated that the glioma prognosis risk could be associated with immune status (Additional file 3: Figure S8). We also explored the correlation of the expression of target genes with cancer stem cell-like properties (RNAss, DNAss) and the TME (stromal score, immune score, and ESTIMATE score). We found that PCG1 was negatively associated with RNAss, the stromal score, the immune score, and the ESTIMATE score. SCAF11 was only negatively associated with DNAss. The rest of the genes showed positive correlations with RNAss, DNAss and the stromal, immune and ESTIMATE scores (Additional file 3: Figure S9).

### Molecular alterations of pyroptosis-related genes based on the prognostic signature

Molecular alterations of pyroptosis-related genes were also evaluated based on histology in the TCGA dataset. NLRP2, NLRP7, and PLCG1 were the only gene alterations in LGG, and NLRP3, NLRP7, NLRP2, SCAF11, NOD1, PLCG1, NLRP1, and CASP1 were gene alterations in GBM. All gene alterations were within 2% (Figure 7). The somatic copy number alteration analysis indicated significant differences among the pyroptosis-related genes. Among these genes, the copy variation number was significantly increased in GPX4, NLRP7, NLRP2, CASP3, CASP6, IL1B, CASP8, IL6, AIM2, NLRP4, NLRP3, PRKACA, ELANE, SCAF11, CASP9, NOD1, and PLCG1 and was significantly decreased in GSDMB, GSDMD, NLRP1, CASP9, TIRAP, CASP1, CASP4, NOD2, CASP5, PYCARD, GSMDC, GSMDA, and IL18 in the high-risk group. The DNA methylation levels of the pyroptosis-related genes were also compared. The results showed that the overall DNA methylation levels were significantly decreased in the high-risk group and increased in the low-risk group.

**Figure 7.**
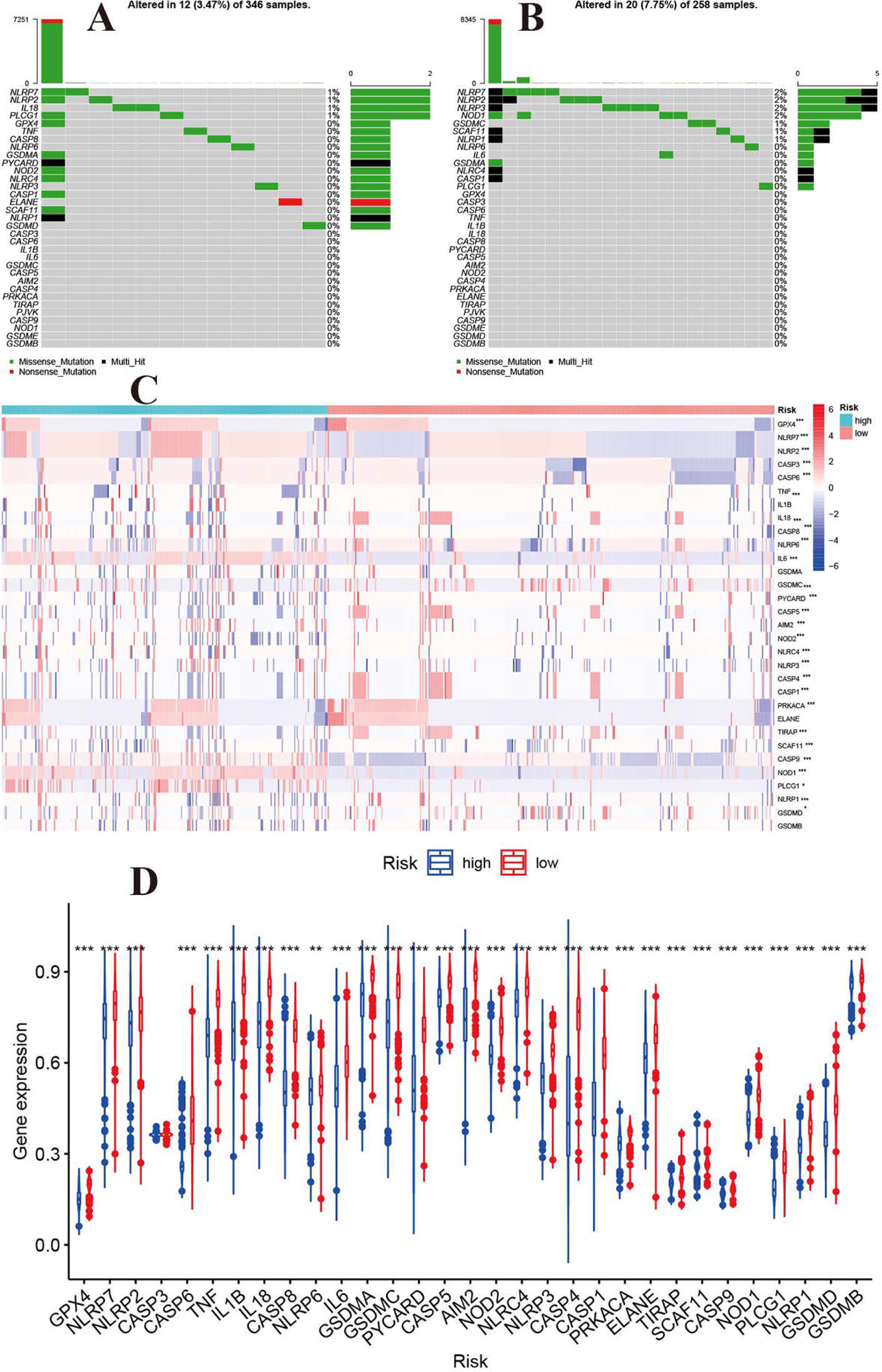
Molecular alterations of pyroptosis-related genes in TCGA dataset. **(A)** The mutations frequencies in low-risk group. **(B)** The mutations frequencies in high-risk group. **(C)** Somatic copy number alteration based on risk groups. **(D)** DNA methylation expression based on risk groups.

### Construction of a ceRNA network based on the prognostic signature

A ceRNA network was constructed based on the differentially expressed mRNAs, lncRNAs and miRNAs between the high-risk and low-risk groups in the TCGA dataset. We identified 763 downregulated mRNAs, 1176 upregulated mRNAs, 116 downregulated lncRNAs, 132 upregulated lncRNAs (Additional file 2: Table S10), 47 downregulated miRNAs and 71 upregulated miRNAs (Additional file 2: Table S11). Finally, 39 mRNAs (28 upregulated and 11 downregulated), 26 lncRNAs (15 upregulated and 15 downregulated) and 14 miRNAs (13 upregulated and 1 downregulated) were included in the ceRNA network (Figure 8). The Kaplan-Meier curves suggested that 13 lncRNAs (positive correlation: AC025211.1, AC068643.1, GDNF-AS1, and LINC00519; negative correlation: ADH1L1-AS2, CRNDE, FAM181A-AS1, HOTAIRM1, MCF2L-AS1, MIR210HG, NEAT1, SLC6A1, and SNHG9; Additional file 3: Figure S10), 41 mRNAs (Additional file 2: Table S12 and Additional file 3: Figure S11) and 8 miRNAs (miR-21, miR-155, miR-200a, miR-216a, miR-221, miR-222, miR-429, and miR-503; Additional file 3: Figure S12) were associated with OS in glioma patients.

**Figure 8.**
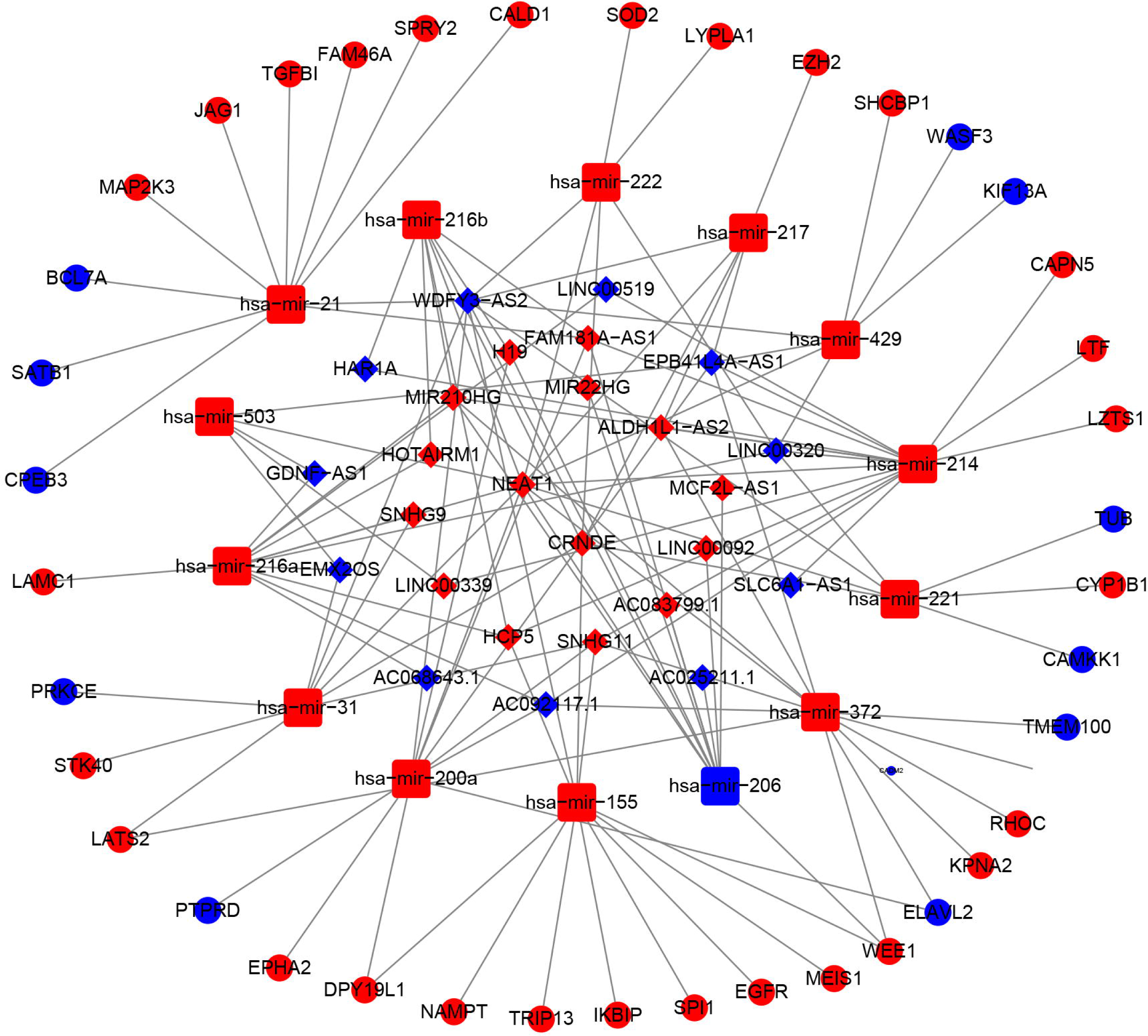
The ceRNA network based on risk groups in TCGA dataset (red: up-regulation. blue: down-regulation).

### Drug sensitivity analysis

To identify potential target drugs, we performed correlations of the identified prognostic signature genes with drugs. We identified 257 pairs of significant gene-drug correlations (Additional file 2: Table S13). There were 9 pairs with correlation coefficients >0.5 or <-0.5.

ELANE-hydroxyurea, ELANE-cyclophosphamide, CASP3-nelarabine, NOD2-imiquimod, NLRP3-rebimastat, ELANE-ABT-199, ELANE-imexon, and NOD2-isotretinoin showed drug sensitivity. PRKACA-cobimetinib showed drug resistance (Figure 9).

**Figure 9.**
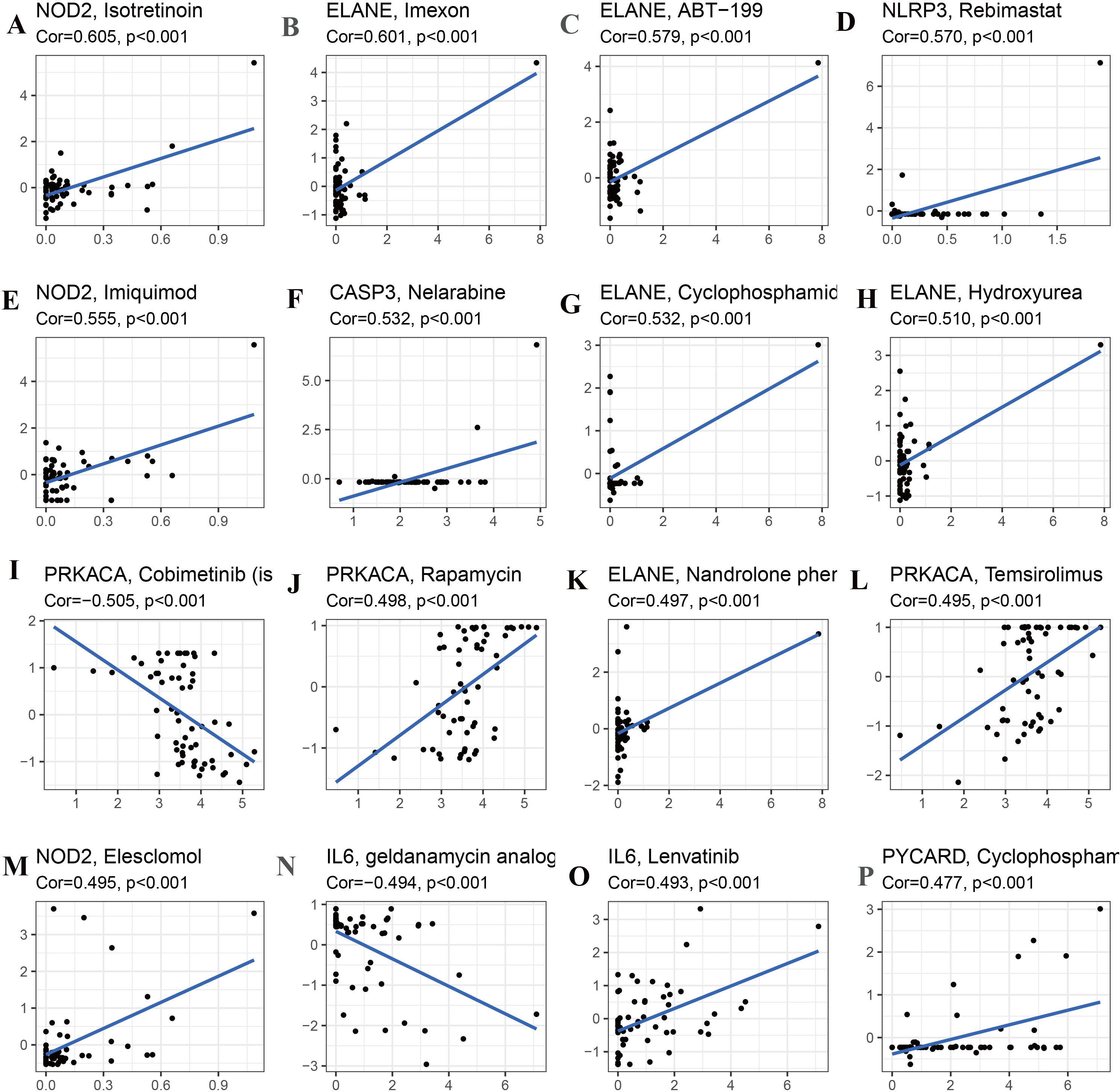
Drug sensitivity analysis for identified prognostic-related genes based on TCGA dataset (Top 16). **(A)** NOD2 and isotretinoin. **(B)** ELANE and Imexon. **(C)** ELANE and ABT-199. **(D)** NLRP3 and Rebimastat. **(E)** NOD2 and Imuiquimod. **(F)** CASP3 and Nelarabine. **(G)** ELANE and Cyclophosphamid. **(H)** ELANE and Hydroxyurea. **(I)** PRKACA and Cobimetinib. **(J)** PRKACA and Rapamycin. **(K)** ELANE and Nandrolone. **(L)** PRKACA and Temsirolimus. **(M)** NOD2 and Eleschomol. **(N)** IL6 and geldanamycin. **(O)** IL6 and Lenvatinib. **(P)** PYCARD and Cyclophospharr

### CASP8 promotes the progression of glioma cells

We selected the CASP8 to validate the molecular function because CASP8 showed significant differences between normal tissue and GBM or LGG in GTEx database (Additional file 3: Figure S13). We firstly detected the expression of CASP8 in glioma cell lines using the Western blot analysis, and found CASP8 is the most highly expressed in LN299 cell. We built the CASP8-si LN229, H4 and U87 cells of glioma. The qPCR indicated mRNA level of CASP8 is significantly down-regulated in U87 and LN229 cells. Furthermore, the silence of CASP8 expression inhibited the cell migration ability (Figure 10). The clonogenic assay also showed that the number of clonogenicity of U87 and LN229 cells were significantly suppressed after knockout of CASP8. These results suggested that CASP8 promotes the progression of glioma cells.

**Figure 10.**
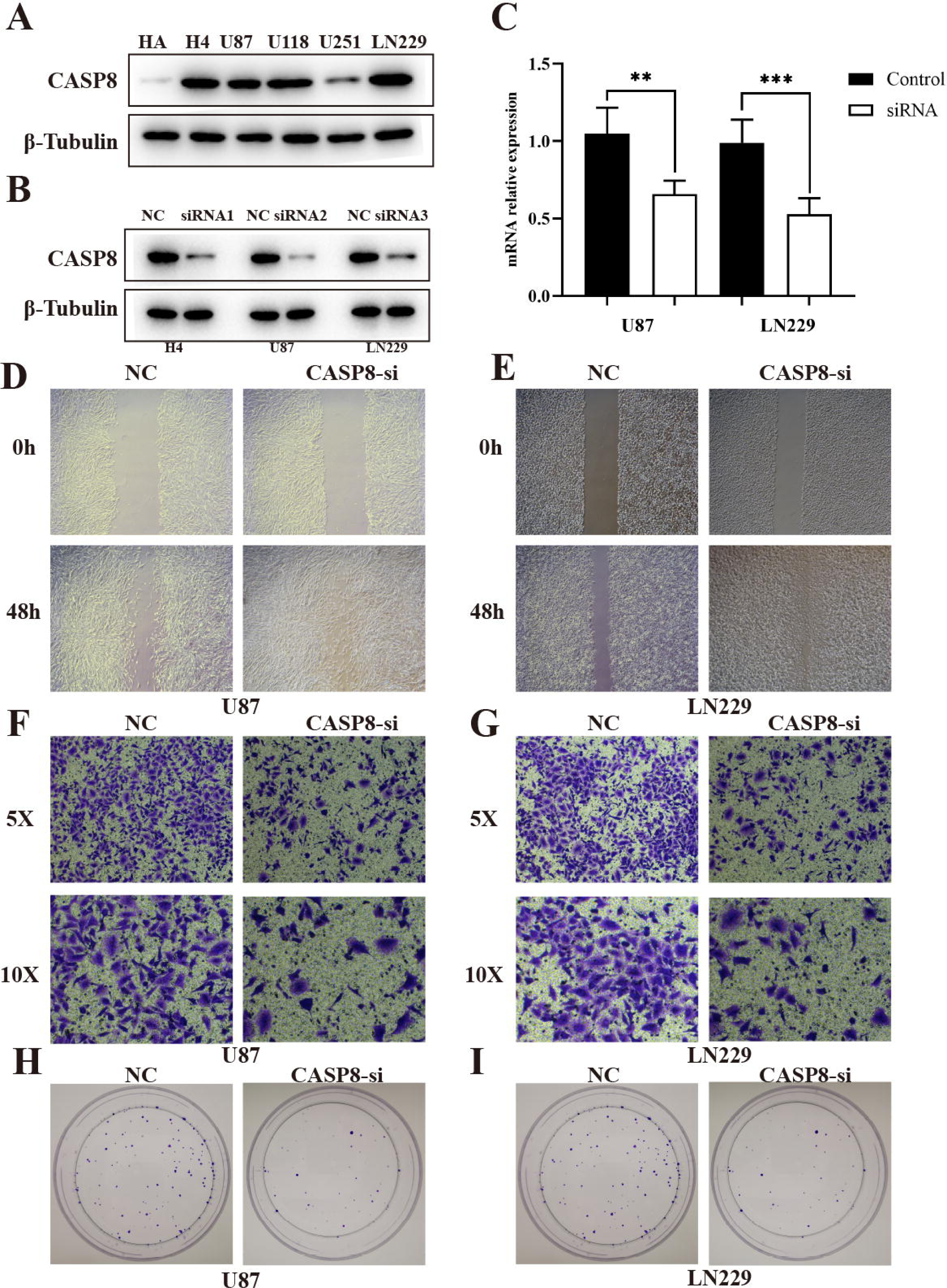
CASP8 promotes progression of glioma cells. **(A)** The expression of CASP8 protein in human HA and glioma cell lines. **(B)** The western blot of CASP8 in U87, U1251, H4 cell lines after siRNA. **(C)** The mRNA expression level of CSAP8 in U87 and U251 after siRNA. **(D and E)** The scratch assay of CASP8-si in U87 and U251 cell lines. **(F and G)** Transwell assay of CASP8-si U87 and U251 cell lines. (H and I) The clonogenic assay of CASP8 in U87 and U251 cell lines.

## DISCUSSION

The traditional histologic-based classification has some limitations, although this classification system has been updated several times over the years and serves clinicians well. One of the primary limitations is interobserver variability(16). A previous study reported that the concordance for reviewing a case is only approximately 50% among different neuropathologists, especially for astrocytic glioma versus oligodendroglioma(17). The development of genomics has allowed us to better understand the differences in prognosis and molecular features and promote effective treatment in glioma subclasses based on molecular features. Using 30 pyroptosis-related genes, we divided glioma patients into two subtypes. Significant overall survival differences were observed between cluster 1 and cluster 2. GSVA indicated that cluster 1 was enriched in some immune-related pathways. Cluster 1 and cluster 2 showed absolute differences in immune cells and immune functions. The infiltration levels of all kinds of immune cells, except Th1 cells, were higher in cluster 1, which had a poor prognosis than in cluster 2. Cluster 1 also showed more significant trends in some main immune function levels, such as immune checkpoints, inflammation promotion, par inflammation. A recent study reported that pyroptosis presents antitumor immune function in tumors, namely, pyroptosis-induced inflammation triggers robust antitumor immunity and can synergize with checkpoint blockade(18). Moreover, some key pathways were also highly enriched in cluster 1, such as the NOD-like receptor signaling pathway, Toll-like receptor signaling pathway, and cytosolic DNA sensing pathway, which were reported to be involved in glioma progression(19-21). These results indicated that pyroptosis-related genes divided glioma patients into two-dimensional distributions well.

We established a prognostic signature based on 15 pyroptosis-related genes. This prognostic signature was well validated in an external independent cohort, and in terms of its predictability, AUCs of 0.844, 0.863, and 0.874 were achieved for 1, 2, and 3 years, respectively, which showed its high discernibility. Combining clinical features and the risk score of the 15 genes, we developed a nomogram for clinical application. The CGGA and TCGA datasets showed high consistency. These results indicated that the prognostic signature based on pyroptosis-related genes has high clinical value.

The signature genes were involved in two biological mechanisms of pyroptosis. The assembly of inflammasome bodies is the initial step of the classical pyroptosis pathway. The inflammasome is a macromolecular protein complex in the cytoplasm necessary for the occurrence of inflammation and can recognize dangerous signaling molecules such as bacteria and viruses. The inflammasome is mainly composed of pattern recognition receptors (PRRs), apoptosis-associated speck-like protein (ASC) and pro-caspase-1 precursors(22). PRRs are receptor proteins responsible for recognizing different signal stimuli in cells. They are mainly composed of nucleotide-binding oligomerization domain-like receptor protein (NLRP) 1, NLRP3, nucleotide-binding oligomerization domain-like receptor protein C4 (NLRC4), absent in melanoma 2 (AIM2) and other components(15). ASC is an adaptor protein that is mainly composed of the N-terminal pyrindomain (PYD) and the C-terminal caspase activation and recruitment domain (CARD)(23). Procaspase-1 is an effector molecule that can specifically cleave GSDMD after activation. After the danger signal sensor NLR1, NLRP3 or AIM2 recognizes the danger signal molecule, the N-terminal PYD is combined with the N-terminal PYD of the adaptor protein. ASC then recruits Caspase-1 through the interaction of the CARDCARD domain to complete the assembly of the inflamed body(24). This method of cell death mediated by Caspase-1 is called the classical pathway of pyroptosis(25). The non-classical pathway of pyrolysis is mainly mediated by Caspase-4, Caspase-5 and Caspase-11. After cells are stimulated by bacterial LPS, Caspases-4, -5, and -11 directly bind to bacterial LPS and are activated(26). Activated Caspases-4, -5, and -11 specifically cleave GSDMD and release the intramolecular inhibition of the GSDMD-N domain(27). The combination of the GSDMD-N-terminus and cell membrane phospholipids causes cell membrane pore formation, cell swelling and rupture and induces cell pyrolysis; the GSDMD-N-terminus can also activate Caspase-1 by activating the NLRP3 inflammasome(28). Activated Caspase-1 stimulates the maturation of IL-18 and IL-1β precursors, and IL-18 and IL-1β are secreted to the outside of the cell and amplify the inflammatory response. Yang et al found that in the nonclassical pathway that relies on Caspase-11, gap junction protein-1 (Pannexin-1) can be cleaved, and the cleavage of Pannexin-1 can activate its own channel and release ATP, which induces pyrolysis(29). Lamkanfi et al found that in the nonclassical pathway that relies on Caspase-11, Pannexin-1 cleavage can also activate the NLRP3 inflammasome, which in turn activates Caspase-1 and induces the occurrence of pyroptosis(30). According to the results, mutations of pyroptosis-related genes are mainly attributed to the classical pathway of pyrolysis. More research is needed to validate the molecular mechanisms.

Based on the risk score, we classified glioma patients into high- and low-risk groups to discriminate clinical outcomes. We further explored the molecular features between the high- and low-risk groups. The functional enrichment analysis results were similar in the TCGA and CGGA datasets, and the same pathways appeared in the two datasets, such as ECM-receptor interaction, GABAergic synapse, focal adhesion, and extracellular matrix organization. The immune cells and immune functions showed similar trends: immune cell and functional scores were higher in the high-risk group. The clinical features showed that cluster 1 had a higher risk score and poorer prognosis than cluster 2. The results indicated that the classification was accurate and validated in the risk model. Furthermore, we compared the gene alterations, CNVs, and DNA methylation levels. Significantly different levels were observed, which reflected the different molecular features of the different risk groups. The ceRNA network identified several key lncRNA-miRNA-mRNA regulatory networks: FAM181A-AS1-miR-21- (CPEB3, SAIB1, BLC7A, MAP2K3, JAG1, TGFBI, FAM46A, SPRY2, and CALD1).

The survival analysis further suggested the regulatory correlation: elevated FAM18A-AS1 and miR-21 were associated with poor prognosis in glioma, and low expression of BCL7A, SATB1 and CPEB3 was associated with favorable prognosis. Previous experiments have reported the promoting role of miR-21 in glioma(31), and upregulation of SATB1 and CPEB3 is associated with the development and progression of glioma(32, 33). The drug sensitivity analysis indicated that NOD2, ELANE, CASP3, and PYCARD showed sensitivity to small molecular drugs, and PRKACA, IL6, and NLLRP3 showed resistance to some drugs. It was reported that the inhibition of the NLRP3 inflammasome by beta-hydroxybutyrate can suppress the migration of glioma cells(34). These results may provide some guidelines for clinical practice.

The present study indicated that pyroptosis-related genes can be used to classify glioma patients into two subclasses based on different molecular features and clinical characteristics. The established prognostic model based on 15 pyroptosis-related genes not only predicted the prognosis of glioma patients but also reflected the molecular alterations, immune infiltration statuses, and stem cell-like properties of different risk groups. The classification based on the risk score of prognostic signature genes revealed a lncRNA-miRNA-mRNA regulatory network. The correlation of signature genes with drug sensitivity may provide a rationale for clinical applications. Finally, our study provides a new understanding of pyroptosis in the development and progression of glioma and contributes new important insights for promoting glioma treatment strategies.

### Authors’ contributions

ZZL designed this study and directed the research group in all aspects, including planning, execution, and analysis of the study. LS drafted the manuscript. YYL, NL, YJZ, QZ collected the data. LZZ provided the statistical software, performed the data analysis, YYL arranged the Figures and Tables. SLF revised the manuscript. All authors have read and approved the final version of the manuscript.

## Funding

This study was supported by the National Natural Science Foundation of China (No. 82003239), Hunan Province Natural Science Foundation (Youth Foundation Project) (NO.2019JJ50945), and the Science Foundation of Xiangya Hospital for Young Scholar (NO. 2018Q012).

## Conflict of Interest Statement

The authors declare that they have no competing interests

## Data Availability Statements

All data can be download from TCGA database (https://portal.gdc.cancer.gov/) and CGGA (http://www.cgga.org.cn/)

## Supplementary materials legends

**Additional file 1:**The details of experiments process in vitro

**Additional file 2: Table S1-S12.xlsx**

**Table S1** The 30 pyroptosis associated genes used for classification

**Table S2** Glioma classification pattern

**Table S3** GSVA enrichment analysis between these distinct pyroptosis-regulated clusters

**Table S4** The result of differential expression analysis (Cluster 2 vs Cluster 1)

**Table S5** Functional enrichment analyses of subclass differentially expressed genes (Cluster 2 vs Cluster 1)

**Table S6** Pathway enrichment analysis of differentially expressed genes from two subclasses

**Table S7** 15 identified pyroptosis-related signature genes in prognostic model

**Table S8** Differentially expressed genes from CGGA based on risk score

**Table S9** Differentially expressed genes from TCGA based on risk score

**Table S10** Differentially expressed lncRNA from TCGA based on risk score

**Table S11** Differentially expressed mir-RNA from TCGA based on risk score

**Table S12** Prognosis-related genes in the ceRNA network

**Table S13** Results of drug sensitivity based on 15 pyroptosis-related prognostic signature genes

## Additional file 3

**Figure S1** Comparisons of different clinical parameters for pyroptosis-related genes. **(A)** LGG and GBM. **(B)** WHO II vs WHOIII vs WHO IV. **(C)** IDH: mutations vs wildtyp. **(D)**1p19_status: codel vs non-codel.

**Figure S2** Barplot of GO enrichment analysis for differentially expressed genes based on subclasses.

**Figure S3** KEEG pathways analysis for differentially expressed genes based on subclasses.

**Figure S4** Correlation of glioma subclasses with immune infiltration. **(A)** Immune cells. **(B)** immune function

**Figure S5** Identification of 215 genes risk signature for OS by LASSO regression in the CGGA cohort. (A) Forest plot of univariate cox regression of OS for 30 pyroptosis-related genes. (B) Cross-validation for tuning parameters selection int the LASSO regression. (C) LASSO regression of the 15 OS-related genes. (D) Heatmap showed the association between risk group and clinical parameters and differentially expressed genes of high- and low-risk group.

**Figure S6** Boxplot of risk score among different clinical characteristics. (A) Subclasses: Cluster 1 vs Cluster 2. (B) Outcomes: Dead vs Alive. (C) Histology: GBM vs LGG. (D) Grade: WHO II vs WHO III vs WHO IV. (E) 1p19q status: Codeletion vs non-codeletion. (F) IDH status: Mutant wildtype.

**Figure S7** Subgroup analysis of OS based on risk score. **(A)** LGG. **(B)** GBM. **(C)** IDH wildtype. **(D)** IDH mutation. **(E)** 1p9ql non-codel. **(F)** 1p9ql codel. **(G)** WHO II. **(H)** WHO III. **(I)** WHO IV

**Figure S8** Comparisons of 15 signature genes among different immune subtype.

**Figure S9** Correlation of expression of 15 signature genes with cancer stem cell-like properties (RNAss, DNAss) and TME (Stromal score, Immune score, and ESTIMATE Score. (A) RNAss. (B) DNAss. (C) Stromal score. (D) Immune score. (E) ESTIMATE score.

**Figure S10** Kaplan-**Meier** curves of lncRNAs for OS in the ceRNA network. **(A)** AC025211.1. **(B)** AC068643.1. **(C)**ADH1L1-AS2. **(D)** CRNDE. **(E)** FAM181A-AS1. **(F)** GDNF-AS1. **(G)** HOTAIRM1. **(H)** LINC00519 **(I)** MCF2L-AS1. **(J)** MIR210HG. **(K)** NEAT1 **(L)**SLC6A1. **(M)** SNHG9.

**Figure S11** Forest plot of mRNAs for OS in the ceRNA network

**Figure S12** Kaplan-**Meier** curves of mir-RNAs for OS in the ceRNA network. **(A)** mir-21. **(B)** mir-155. **(C)** mir-200a. **(D)** mir-216a. **(E)** mir-221. **(F)** mir-222. **(G)** mir-429. **(H)** mir-503.

**Figure S13** The expression levels of identified prognostic genes between tumor and normal

## Notes

http://www.cgga.org.cn/

https://portal.gdc.cancer.gov/

### Competing Interest Statement

The authors have declared no competing interest.

