## Supplementary material Figure S1-S16 for "Comprehensive analysis of pyroptosis-associated in molecular classification, immunity and prognostic of glioma"

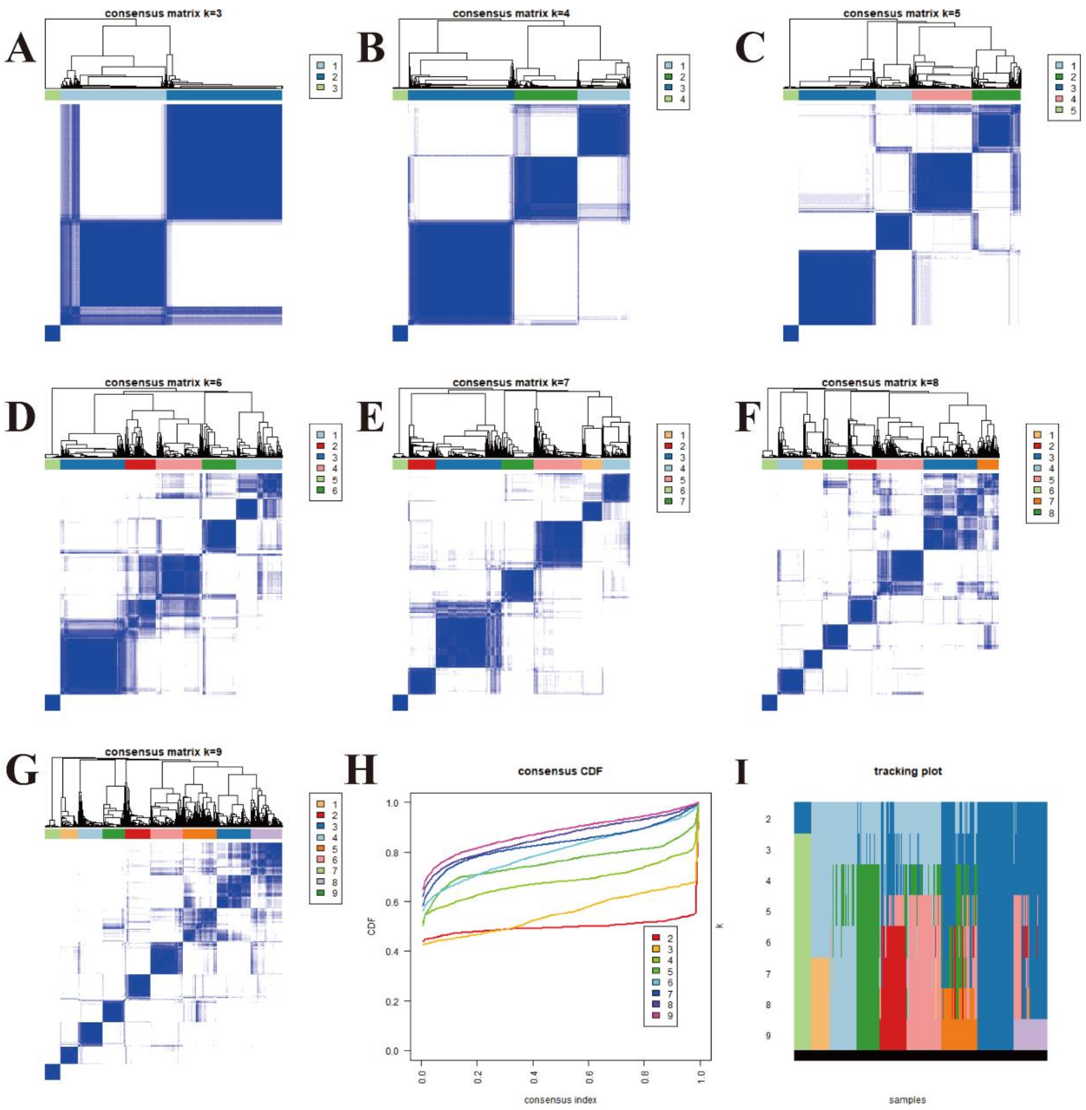
A
consensus matrix k=3
B
consensus matrix k=4
C
consensus matrix k=5
D
consensus matrix k=6
E
consensus matrix k=7
F
consensus matrix k=8
G
consensus matrix k=9
H
consensus CDF
CDF
consensus index
I
tracking plot
k
samples

Figure S1

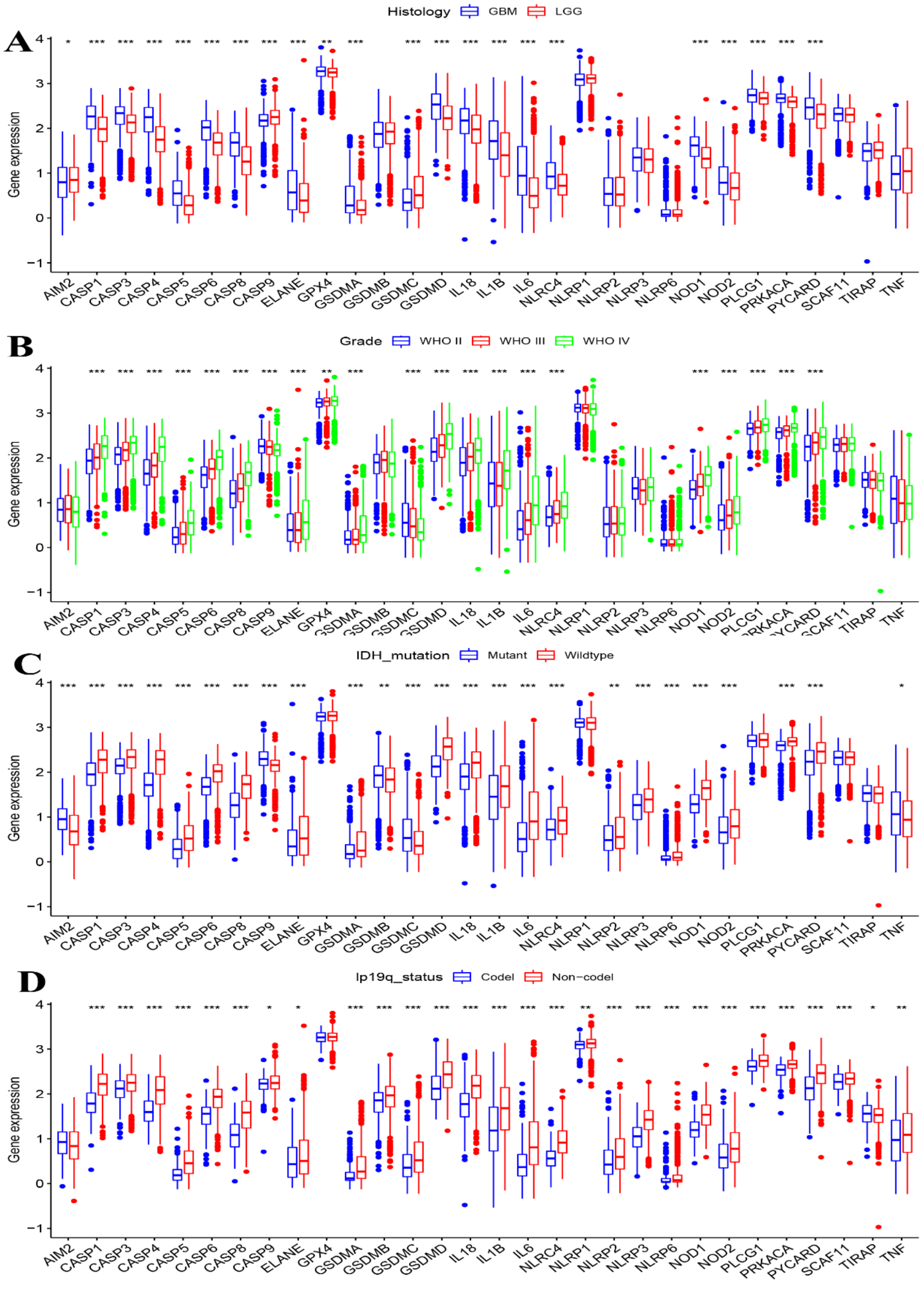
A
Histology GBM LGG
B
Grade WHO II WHO III WHO IV
C
IDH_mutation Mutant Wildtype
D
Ip19q_status Codel Non-codel
Gene expression
AIM2 CASP1 CASP3 CASP4 CASP5 CASP6 CASP8 CASP9 ELANE GPX4 GSDMA GSDMB GSDMC GSDMD IL18 IL1B IL6 NLRC4 NLRP1 NLRP2 NLRP3 NLRP6 NOD1 NOD2 PLCG1 PRKACA PYCARD SCAF11 TIRAP TNF

Figure S2

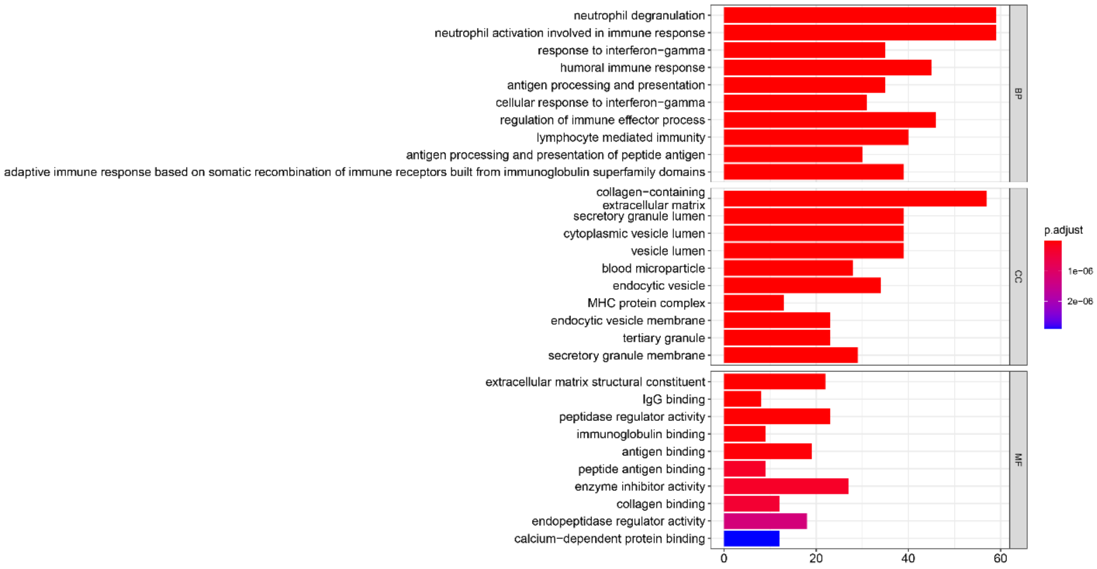


Figure S3

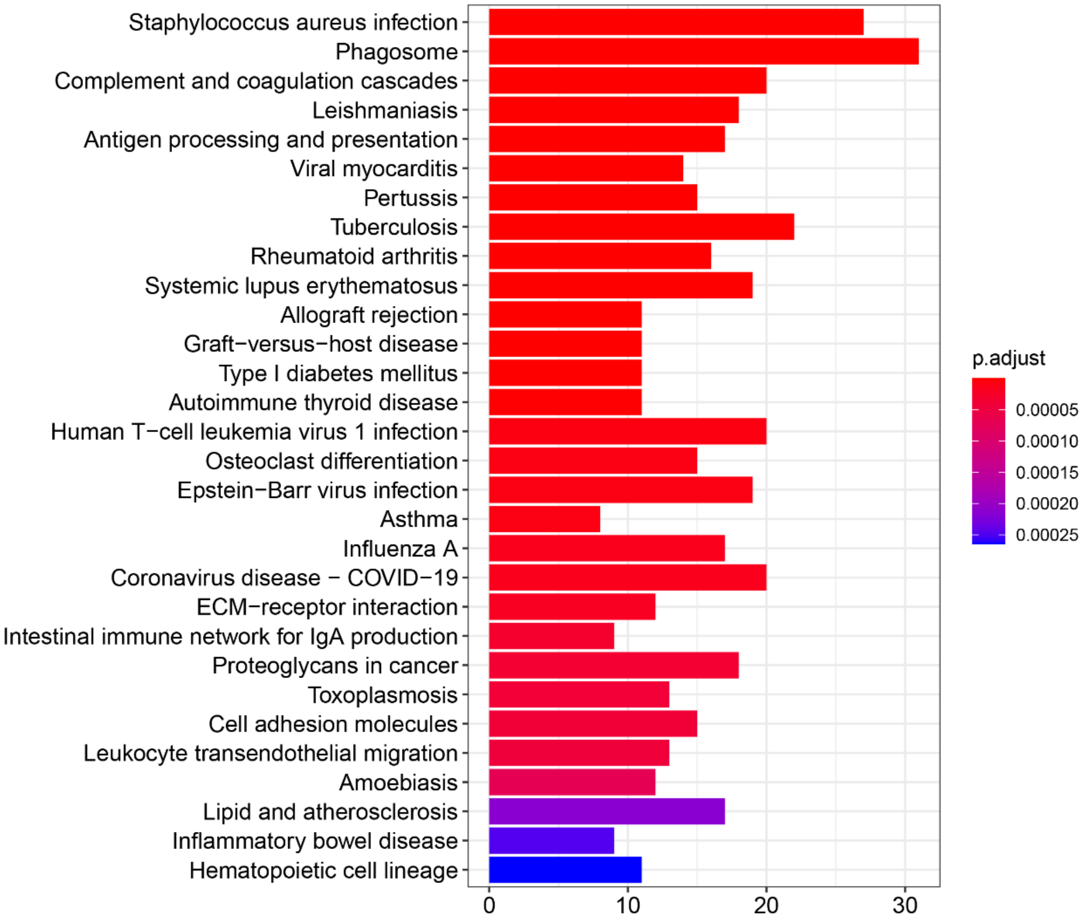


Figure S4

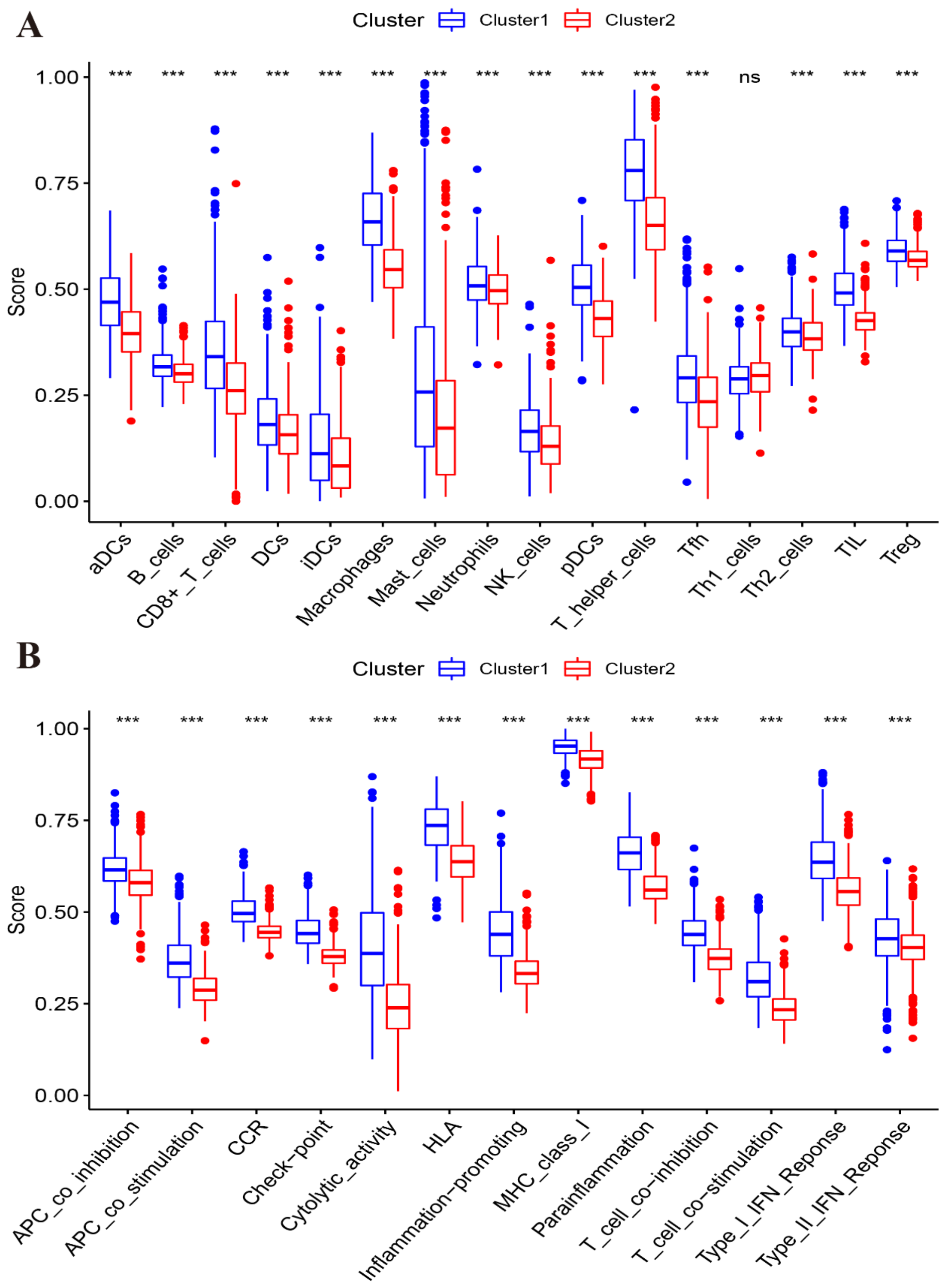
A
Cluster
Cluster1
Cluster2
Score
1.00
0.75
0.50
0.25
0.00
*** *** *** *** *** *** *** *** *** *** *** *** ns *** *** ***
aDCs
B_cells
CD8+_T_cells
DCs
iDCs
Macrophages
Mast_cells
Neutrophils
NK_cells
pDCs
T_helper_cells
Tfh
Th1_cells
Th2_cells
TIL
Treg
B
Cluster
Cluster1
Cluster2
Score
1.00
0.75
0.50
0.25
0.00
*** *** *** *** *** *** *** *** *** *** *** *** ***
APC_co_inhibition
APC_co_stimulation
CCR
Check-point
Cytolytic_activity
HLA
Inflammation-promoting
MHC_class_I
Parainflammation
T_cell_co-inhibition
T_cell_co-stimulation
Type_I_IFN_Reponse
Type_II_IFN_Reponse

Figure S5

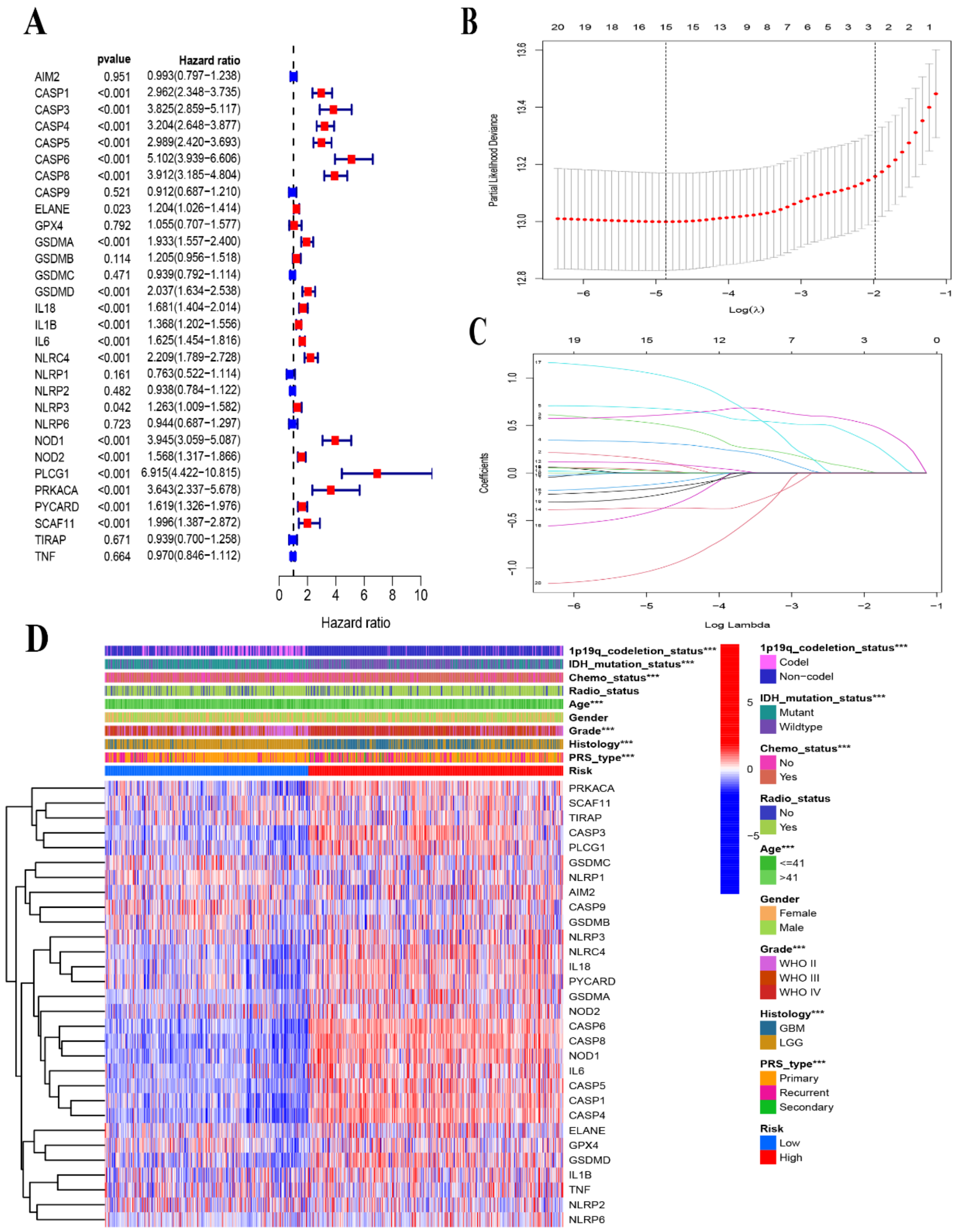

A
pvalue
Hazard ratio
AIM2 0.951 0.993(0.797−1.238)
CASP1 <0.001 2.962(2.348−3.735)
CASP3 <0.001 3.825(2.859−5.117)
CASP4 <0.001 3.204(2.648−3.877)
CASP5 <0.001 2.989(2.420−3.693)
CASP6 <0.001 5.102(3.939−6.606)
CASP8 <0.001 3.912(3.185−4.804)
CASP9 0.521 0.912(0.687−1.210)
ELANE 0.023 1.204(1.026−1.414)
GPX4 0.792 1.055(0.707−1.577)
GSDMA <0.001 1.933(1.557−2.400)
GSDMB 0.114 1.205(0.956−1.518)
GSDMC 0.471 0.939(0.792−1.114)
GSDMD <0.001 2.037(1.634−2.538)
IL18 <0.001 1.681(1.404−2.014)
IL1B <0.001 1.368(1.202−1.556)
IL6 <0.001 1.625(1.454−1.816)
NLRC4 <0.001 2.209(1.789−2.728)
NLRP1 0.161 0.763(0.522−1.114)
NLRP2 0.482 0.938(0.784−1.122)
NLRP3 0.042 1.263(1.009−1.582)
NLRP6 0.723 0.944(0.687−1.297)
NOD1 <0.001 3.945(3.059−5.087)
NOD2 <0.001 1.568(1.317−1.866)
PLCG1 <0.001 6.915(4.422−10.815)
PRKACA <0.001 3.643(2.337−5.678)
PYCARD <0.001 1.619(1.326−1.976)
SCAF11 <0.001 1.996(1.387−2.872)
TIRAP 0.671 0.939(0.700−1.258)
TNF 0.664 0.970(0.846−1.112)
Hazard ratio
B
Partial Likelihood Deviance
Log(λ)
C
Coefficients
Log Lambda
D
1p19q_codeletion_status***
IDH_mutation_status***
Chemo_status***
Radio_status
Age***
Gender
Grade***
Histology***
PRS_type***
Risk
PRKACA
SCAF11
TIRAP
CASP3
PLCG1
GSDMC
NLRP1
AIM2
CASP9
GSDMB
NLRP3
NLRC4
IL18
PYCARD
GSDMA
NOD2
CASP6
CASP8
NOD1
IL6
CASP5
CASP1
CASP4
ELANE
GPX4
GSDMD
IL1B
TNF
NLRP2
NLRP6
1p19q_codeletion_status***
Codel
Non−codel
IDH_mutation_status***
Mutant
Wildtype
Chemo_status***
No
Yes
Radio_status
No
Yes
Age***
<=41
>41
Gender
Female
Male
Grade***
WHO II
WHO III
WHO IV
Histology***
GBM
LGG
PRS_type***
Primary
Recurrent
Secondary
Risk
Low
High


Figure S6

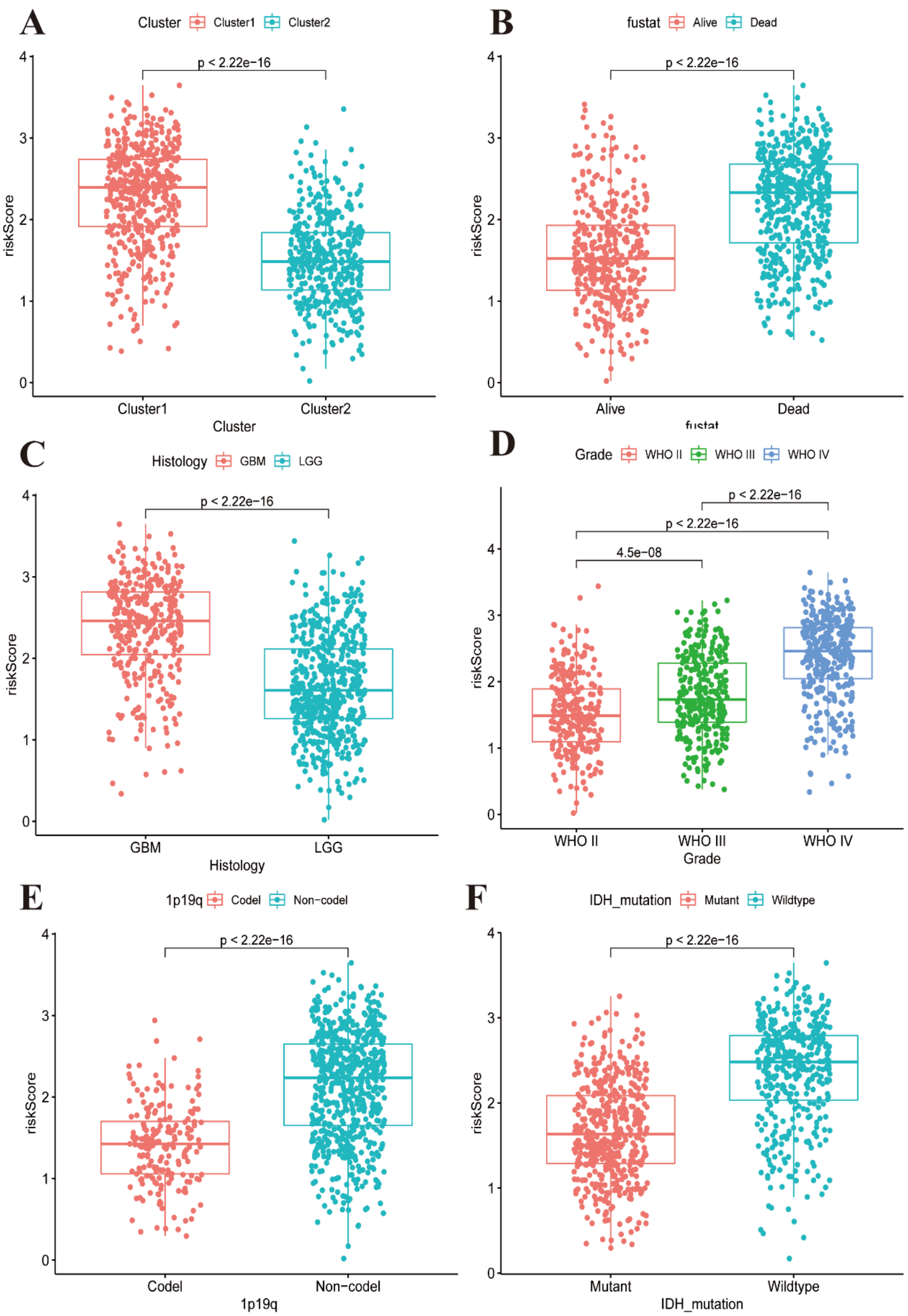
A
Cluster Cluster1 Cluster2
p < 2.22e−16
riskScore
Cluster1 Cluster2
Cluster
B
fustat Alive Dead
p < 2.22e−16
riskScore
Alive Dead
fustat
C
Histology GBM LGG
p < 2.22e−16
riskScore
GBM LGG
Histology
D
Grade WHO II WHO III WHO IV
p < 2.22e−16
p < 2.22e−16
4.5e−08
riskScore
WHO II WHO III WHO IV
Grade
E
1p19q Codel Non−codel
p < 2.22e−16
riskScore
Codel Non−codel
1p19q
F
IDH_mutation Mutant Wildtype
p < 2.22e−16
riskScore
Mutant Wildtype
IDH_mutation

Figure S7

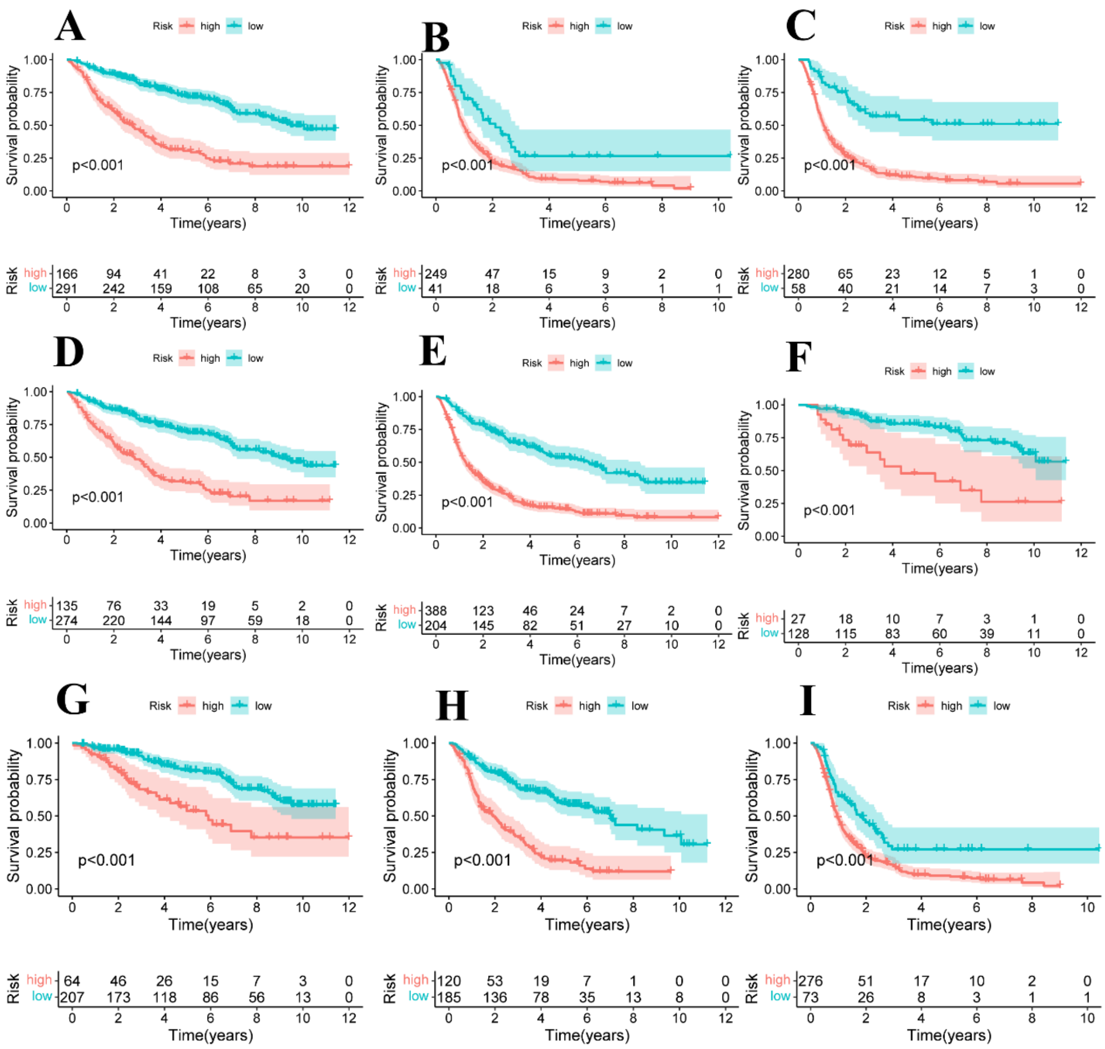
A
B
C
D
E
F
G
H
I
Risk high low
Survival probability
Time(years)
p<0.001
Risk high 166 94 41 22 8 3 0
low 291 242 159 108 65 20 0
Risk high 249 47 15 9 2 0
low 41 18 6 3 1 1
Risk high 280 65 23 12 5 1 0
low 58 40 21 14 7 3 0
Risk high 135 76 33 19 5 2 0
low 274 220 144 97 59 18 0
Risk high 388 123 46 24 7 2 0
low 204 145 82 51 27 10 0
Risk high 27 18 10 7 3 1 0
low 128 115 83 60 39 11 0
Risk high 64 46 26 15 7 3 0
low 207 173 118 86 56 13 0
Risk high 120 53 19 7 1 0 0
low 185 136 78 35 13 8 0
Risk high 276 51 17 10 2 0
low 73 26 8 3 1 1

Figure S8

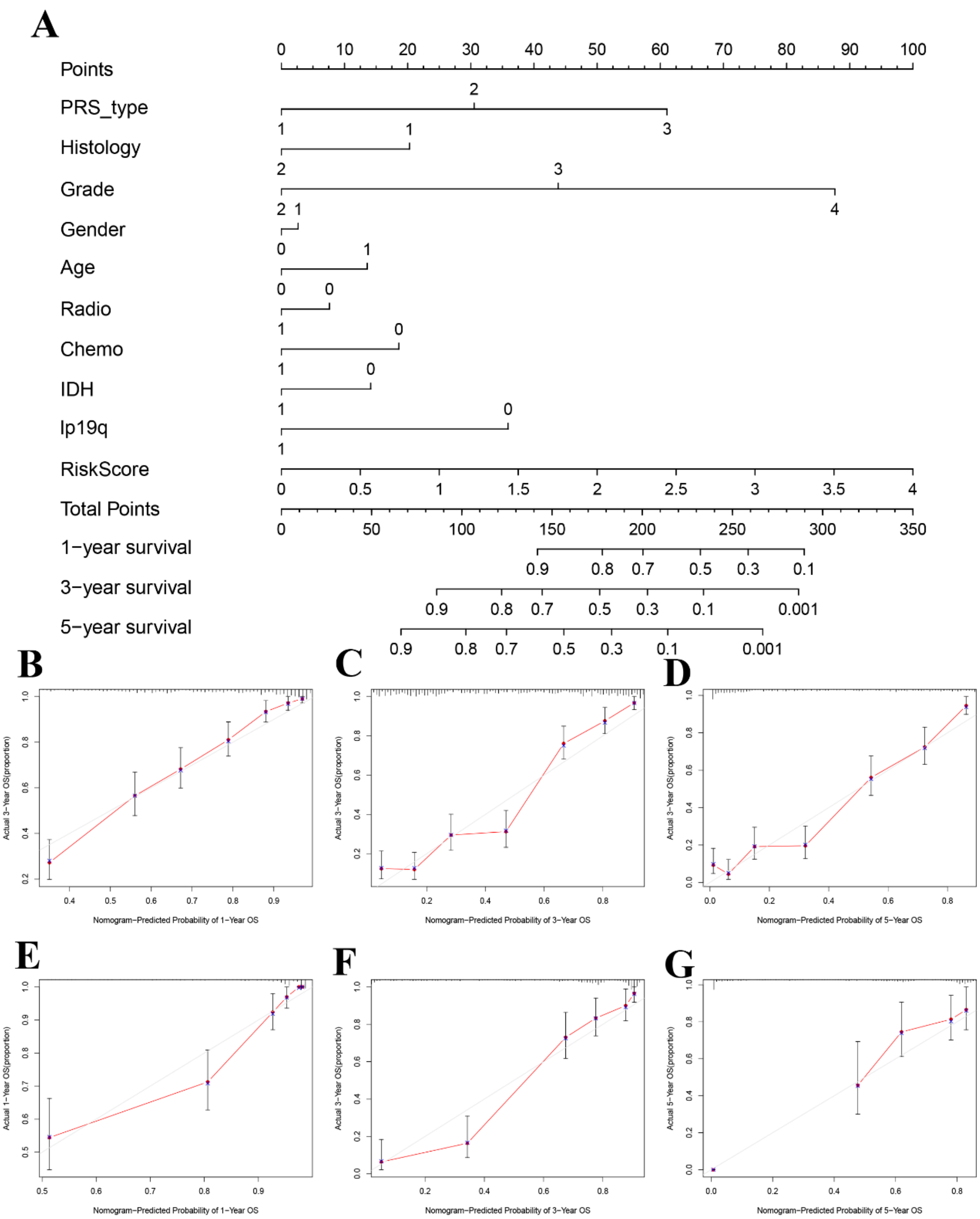
A
Points
PRS_type
Histology
Grade
Gender
Age
Radio
Chemo
IDH
Ip19q
RiskScore
Total Points
1−year survival
3−year survival
5−year survival
B
C
D
E
F
G
Actual 3−Year OS(proportion)
Actual 1−Year OS(proportion)
Actual 5−Year OS(proportion)
Nomogram−Predicted Probability of 1−Year OS
Nomogram−Predicted Probability of 3−Year OS
Nomogram−Predicted Probability of 5−Year OS

Figure S9

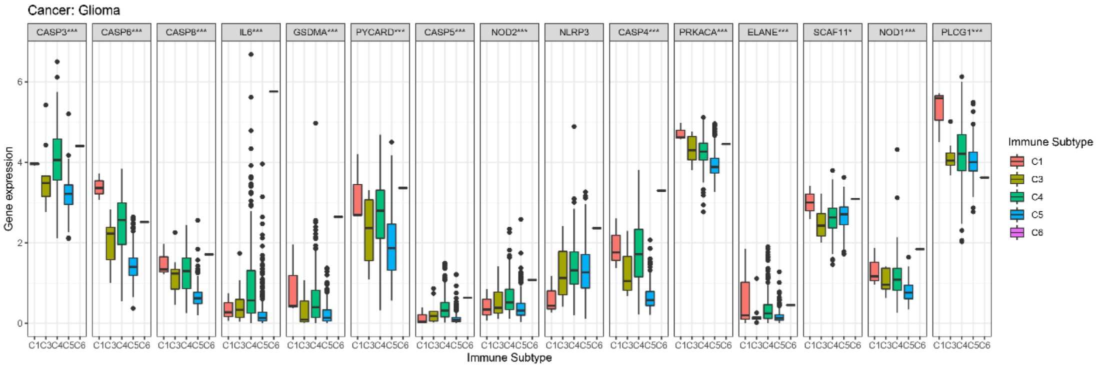


Figure S10

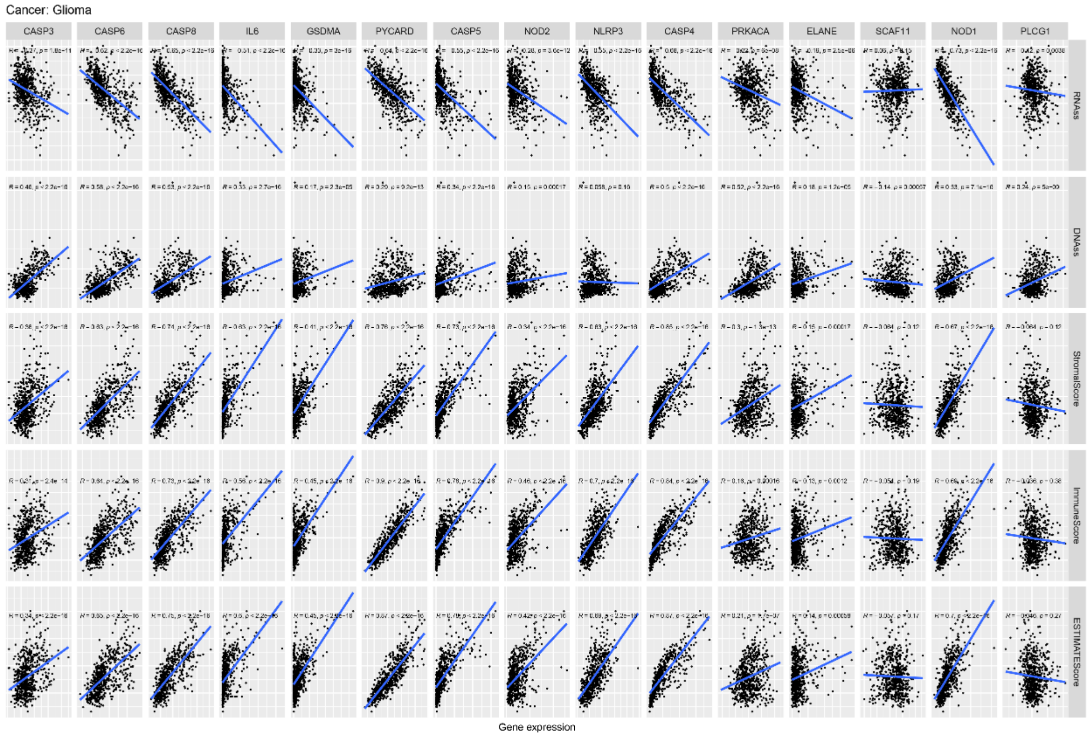


Figure S11

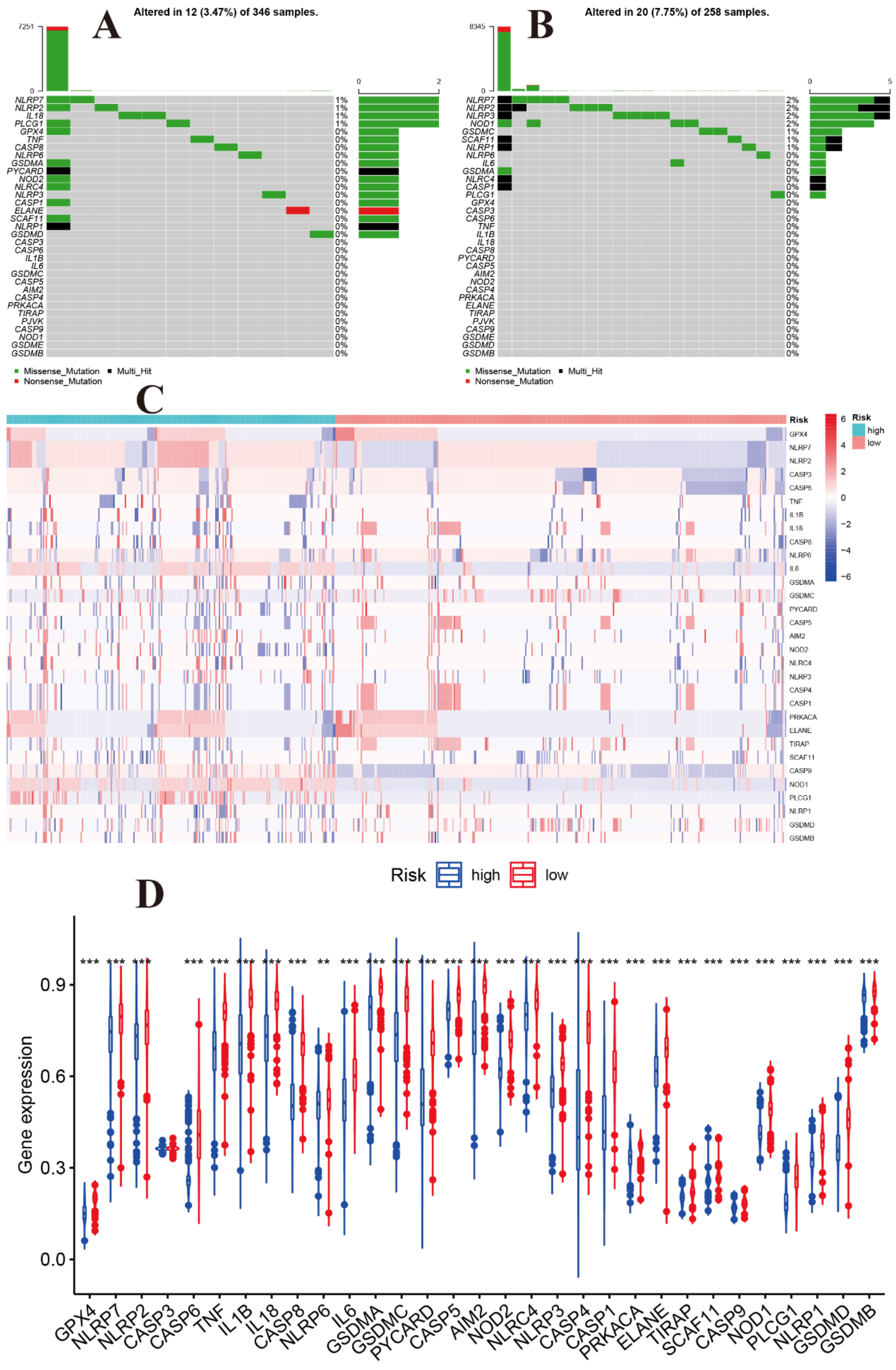
A
Altered in 12 (3.47%) of 346 samples.
B
Altered in 20 (7.75%) of 258 samples.
Missense_Mutation
Multi_Hit
Nonsense_Mutation
C
Risk
high
low
D
Risk
high
low
Gene expression

Figure S12

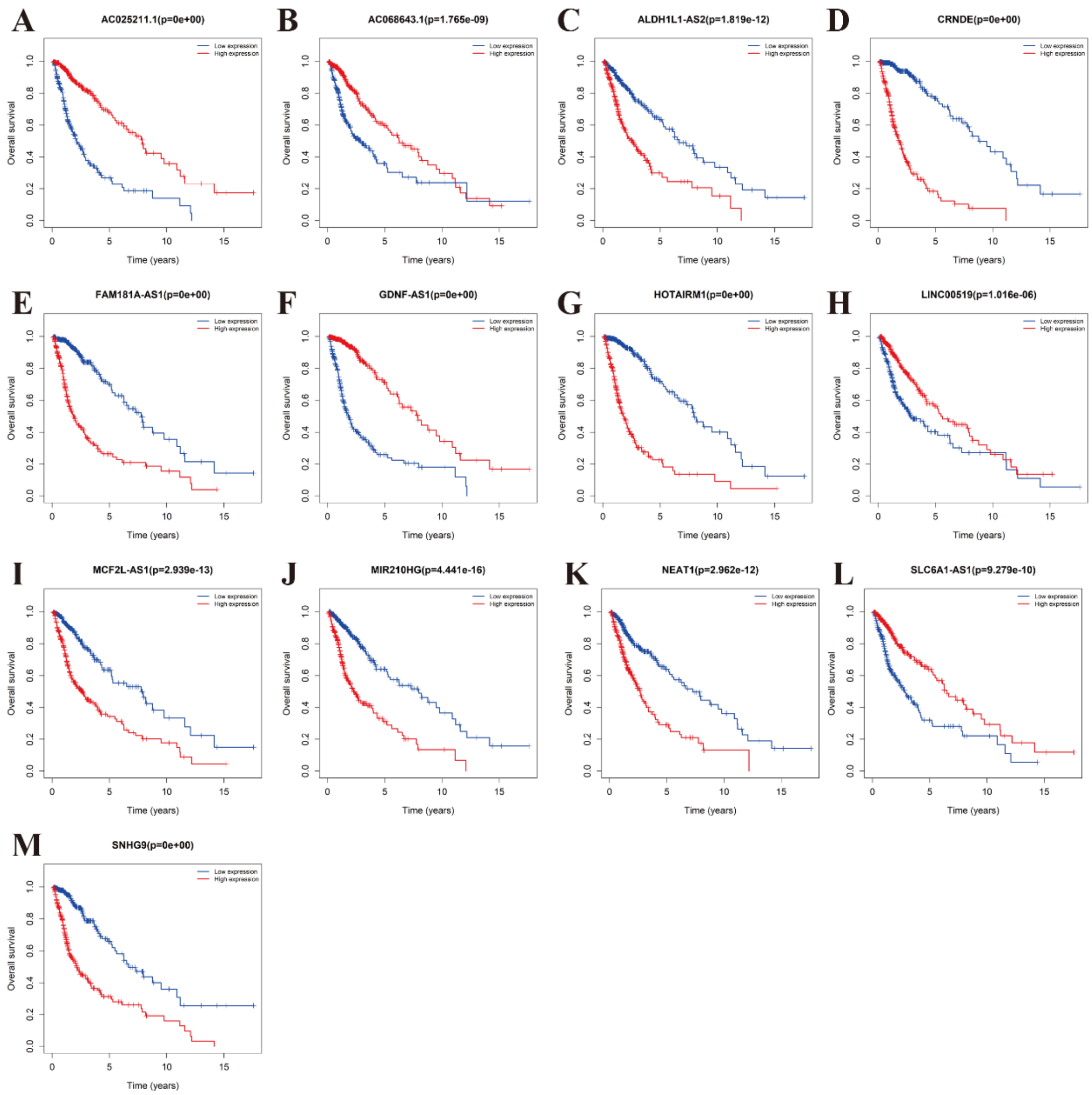
A
AC025211.1(p=0e+00)
B
AC068643.1(p=1.765e-09)
C
ALDH1L1-AS2(p=1.819e-12)
D
CRNDE(p=0e+00)
E
FAM181A-AS1(p=0e+00)
F
GDNF-AS1(p=0e+00)
G
HOTAIRM1(p=0e+00)
H
LINC00519(p=1.016e-06)
I
MCF2L-AS1(p=2.939e-13)
J
MIR210HG(p=4.441e-16)
K
NEAT1(p=2.962e-12)
L
SLC6A1-AS1(p=9.279e-10)
M
SNHG9(p=0e+00)
Low expression
High expression
Overall survival
Time (years)

Figure S13

| | pvalue | Hazard ratio |
|---|---|---|
| EGFR | <0.001 | 1.248(1.145−1.361) |
| EZH2 | <0.001 | 2.056(1.774−2.383) |
| LZTS1 | <0.001 | 1.615(1.443−1.808) |
| PRKCE | <0.001 | 0.290(0.228−0.370) |
| PTPRD | <0.001 | 0.473(0.406−0.550) |
| ELAVL2 | <0.001 | 0.499(0.426−0.585) |
| FAM46A | <0.001 | 2.689(2.343−3.086) |
| SPI1 | <0.001 | 1.719(1.502−1.968) |
| WASF3 | <0.001 | 0.454(0.396−0.520) |
| LYPLA1 | <0.001 | 2.396(2.113−2.716) |
| WEE1 | <0.001 | 2.852(2.486−3.272) |
| TRIP13 | <0.001 | 2.792(2.371−3.287) |
| MAP2K3 | <0.001 | 3.900(3.177−4.787) |
| CPEB3 | <0.001 | 0.210(0.165−0.267) |
| LATS2 | <0.001 | 3.717(3.057−4.519) |
| STK40 | <0.001 | 2.653(2.238−3.146) |
| SPRY2 | <0.001 | 2.376(2.063−2.738) |
| IKBIP | <0.001 | 4.118(3.426−4.951) |
| SHCBP1 | <0.001 | 3.126(2.635−3.709) |
| SOD2 | <0.001 | 1.666(1.535−1.809) |
| CAMKK1 | 0.004 | 0.809(0.701−0.934) |
| CYP1B1 | <0.001 | 1.423(1.247−1.622) |
| BCL7A | <0.001 | 0.385(0.332−0.445) |
| CALD1 | <0.001 | 2.708(2.325−3.153) |
| KPNA2 | <0.001 | 2.940(2.500−3.457) |
| RHOC | <0.001 | 2.219(1.903−2.586) |
| SATB1 | <0.001 | 0.371(0.318−0.432) |
| LTF | <0.001 | 1.289(1.239−1.342) |
| MEIS1 | <0.001 | 2.496(2.085−2.988) |
| KIF13A | <0.001 | 0.272(0.224−0.330) |
| TUB | <0.001 | 0.401(0.351−0.458) |
| DPY19L1 | <0.001 | 2.245(1.991−2.531) |
| JAG1 | <0.001 | 2.015(1.763−2.303) |
| CAPN5 | <0.001 | 1.746(1.554−1.962) |
| TMEM100 | <0.001 | 0.631(0.583−0.683) |
| CADM2 | <0.001 | 0.512(0.460−0.569) |
| TGFBI | <0.001 | 1.557(1.436−1.688) |
| EPHA2 | <0.001 | 2.063(1.806−2.357) |
| NAMPT | <0.001 | 1.890(1.734−2.061) |
| LAMC1 | <0.001 | 2.322(2.055−2.625) |

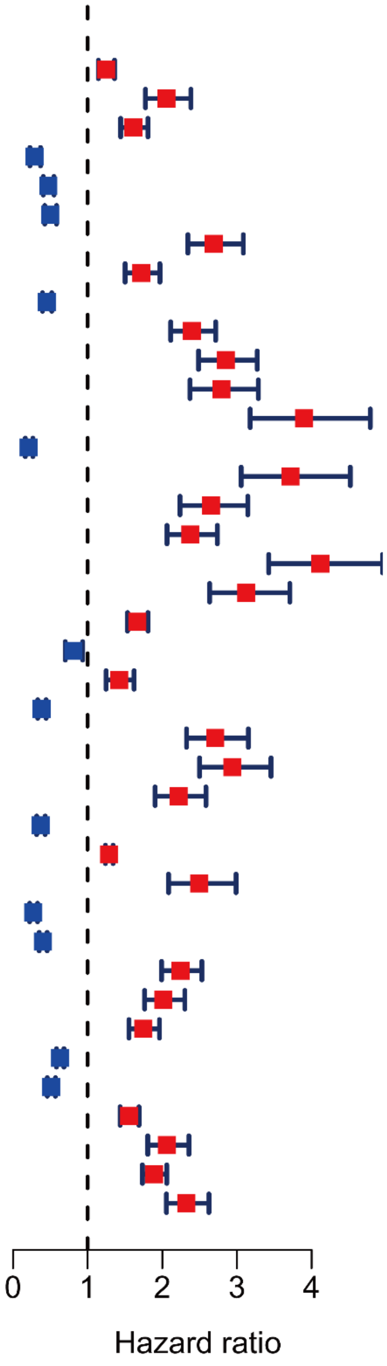


Figure S14

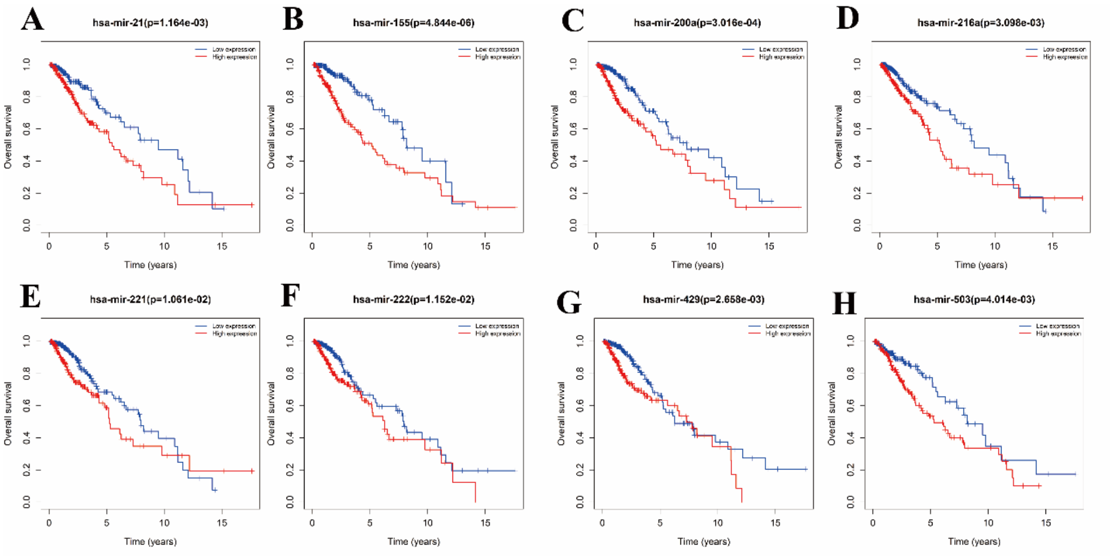
A
hsa-mir-21(p=1.164e-03)
B
hsa-mir-155(p=4.844e-06)
C
hsa-mir-200a(p=3.016e-04)
D
hsa-mir-216a(p=3.098e-03)
E
hsa-mir-221(p=1.061e-02)
F
hsa-mir-222(p=1.152e-02)
G
hsa-mir-429(p=2.658e-03)
H
hsa-mir-503(p=4.014e-03)
Overall survival
Time (years)

Figure S15

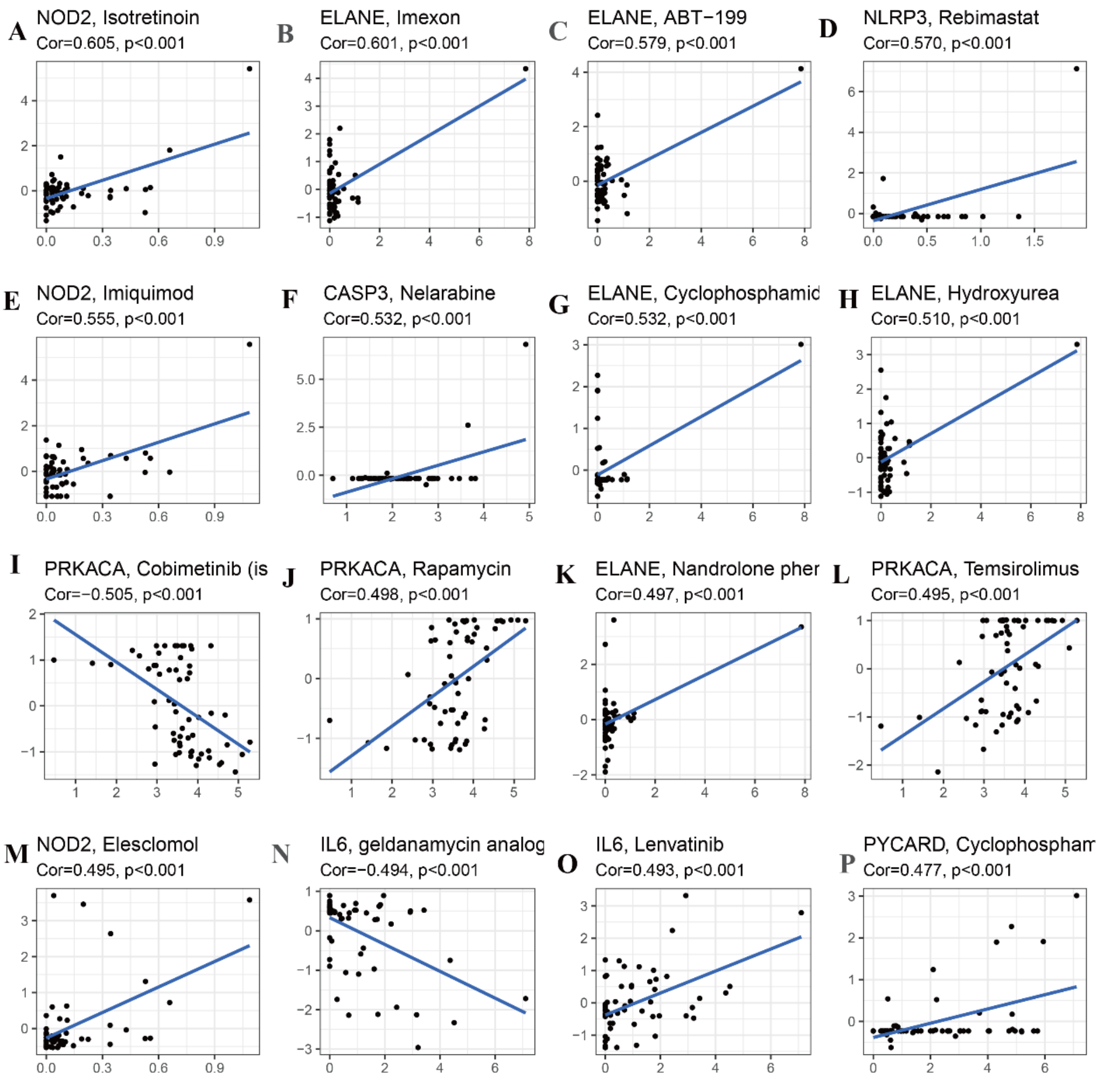
A
NOD2, Isotretinoin
Cor=0.605, p<0.001
B
ELANE, Imexon
Cor=0.601, p<0.001
C
ELANE, ABT−199
Cor=0.579, p<0.001
D
NLRP3, Rebimastat
Cor=0.570, p<0.001
E
NOD2, Imiquimod
Cor=0.555, p<0.001
F
CASP3, Nelarabine
Cor=0.532, p<0.001
G
ELANE, Cyclophosphamid
Cor=0.532, p<0.001
H
ELANE, Hydroxyurea
Cor=0.510, p<0.001
I
PRKACA, Cobimetinib (is
Cor=−0.505, p<0.001
J
PRKACA, Rapamycin
Cor=0.498, p<0.001
K
ELANE, Nandrolone pher
Cor=0.497, p<0.001
L
PRKACA, Temsirolimus
Cor=0.495, p<0.001
M
NOD2, Elesclomol
Cor=0.495, p<0.001
N
IL6, geldanamycin analog
Cor=−0.494, p<0.001
O
IL6, Lenvatinib
Cor=0.493, p<0.001
P
PYCARD, Cyclophospham
Cor=0.477, p<0.001

Figure S16
